# Serial intravascular labeling resolves dynamic decidual NK residency and gestational remodeling toward T-bet-Low states

**DOI:** 10.64898/2026.10.28.762610

**Authors:** Mona A. Mohamed, Jessica Vazquez, Marco A. Cordero, Fernanda Leyva Jaimes, Sophia Beaulieu, Eavan C Finnerty, Xiang Yi Chen, Roghyah Hasan, María Virumbrales-Muñoz, Aleksandar K. Stanic

## Abstract

Decidual natural killer (dNK) cells are the predominant immune population at the maternal-fetal interface during early pregnancy. However, how vascular trafficking, tissue residency, proliferation, and apoptosis-associated activity are coordinated to establish and maintain the dNK population throughout pregnancy remains unclear. We hypothesized that gestational remodeling of the dNK compartment reflects coordinated changes in these cellular processes in association with progressive T-bet downregulation. We combined serial intravascular labeling (SIVL), flow cytometry, BrdU incorporation, active caspase-3 detection, spatial imaging, and *Tbx21* fate mapping to examine NK cells in virgin mice and across gestation. Pregnancy altered CD49a/CD49b-defined dNK-cell composition and redistributed dNK cells among SIVL-defined vascular-accessibility states, supporting a model of dynamic dNK-cell tissue residency. Trafficking-associated marker scores displayed continuous gradients of tissue retention, circulatory/egress potential, and chemokine responsiveness rather than defining a single resident state. Spatial imaging demonstrated greater dNK-cell density in the mesometrial decidua and preferential CD31⁺ vessel contact by recently intravascular dNK cells during early and mid-gestation. BrdU incorporation and active caspase-3 positivity peaked during early pregnancy and varied across SIVL-defined states. Tbx21 fate mapping identified a progressive and selective shift from current T-bet expression toward ex-T-bet and non-fate-mapped Eomes⁺ dNK states. T-bet-low states preferentially maintained proliferation at mid-gestation, whereas current T-bet expression was associated with greater circulatory/egress potential. However, active caspase-3 was not selectively increased in current T-bet-expressing cells. Collectively, these findings identify pregnancy-induced dNK-cell remodeling as a coordinated process integrating dynamic tissue residency, vascular accessibility, local proliferation, and apoptosis-associated activity, alongside a progressive shift toward T-bet-low dNK states.

## Introduction

Decidual natural killer (dNK) cells are the dominant innate lymphoid population at the maternal-fetal interface (MFI), particularly during early pregnancy. They comprise approximately 50-70% of decidual lymphocytes, although their relative abundance declines as gestation progresses^1^. These specialized uterine lymphocytes differ substantially from their peripheral conventional NK (cNK) cell counterparts in phenotype, receptor repertoire, and functional profile^2^. Rather than serving primarily as classical cytotoxic effectors, dNK cells play critical roles in supporting trophoblast invasion and uterine vascular remodeling through the secretion of immunomodulatory factors, including IL-8, VEGF family members, and GM-CSF^3^. However, despite their central role in pregnancy, a major unresolved question remains: which cellular mechanisms regulate dNK abundance and subset composition across gestation? Across both human and mouse studies, the field has debated whether changes in dNK-cell composition are driven predominantly by recruitment from the circulation, local persistence, or phenotypic transition. The central aim of this study was to resolve this question, as similar terminal phenotypes can emerge from distinct underlying cellular programs, leading to different consequences for decidual immune homeostasis and pregnancy outcomes.

Despite a number of studies in the field, the relative contribution of trafficking and local expansion to the dNK cell pool at the MFI is still not entirely resolved^4,5^. There is evidence that circulating cNK cells can differentiate into a dNK-like phenotype in vitro ^6^, actually traffic to MFI ^7^, and that NK-cell precursors^8^ are recruited to the decidua through coordinated chemokine-mediated networks^9–11^. In mice, decidualized uterine grafts from NK-competent donors acquired uNK cells in wild-type but not NK/uNK-deficient hosts, while transferred cells from secondary lymphoid organs of pregnant donors efficiently reconstituted the uNK population in deficient recipients^12^. Consistently, our recent work employed serial intravascular labeling (SIVL) to demonstrate active decidual leukocyte trafficking that proportionally increased across murine gestation. Complementary immunofluorescence imaging in late-gestation cynomolgus macaques detected intravascularly labeled leukocytes within the decidual stroma, supporting recruitment from the circulation into decidual tissue^7^.

Chemokine studies provide additional evidence that trafficking of cNK cells contributes to the formation and maintenance of the dNK-cell pool. In mice, uterine CXCL12 expression was detected at GD0 and GD6, and uNK-cell CXCR4 expression was detected at GD0 and GD12, suggesting a potential role for this axis in NK-cell or precursor recruitment and maturation^13^. Disruption of maternal CXCR4 signaling alters uterine NK-cell distribution and function MFI^10^. Notably, NK cells trafficking to the decidua was found to be independent of CCR2, CCR5, and MIP-1α, indicating that the precise homing mechanisms remain incompletely understood^14^.

In human pregnancy, evidence for dNK-cell trafficking comes primarily from chemokine-receptor and migration studies showing that peripheral blood CD16⁻/CD56^bright^ NK cells are selectively recruited toward decidual tissues through CXCL12-CXCR4 and CXCL9/10-CXCR3 axes, with trophoblasts, decidual stromal cells, and vascular endothelial cells contributing to this chemotactic network^15–18^. Nevertheless, it remains unclear whether these cells are durably retained within the decidua and differentiate into tissue-resident populations after homing, or whether they primarily reflect ongoing cellular trafficking.

On the other hand, evidence showed that the marked accumulation of uNK cells during decidualization is driven predominantly by local proliferation of tissue resident (trNK) cells, classically characterized as CD49a⁺CD49b⁻, rather than by recruitment of circulating cNK cells, which typically exhibit a CD49a⁻CD49b⁺ phenotype^5^. Notably, studies including uterine group 1 innate lymphoid cells (ILC1) also show heterogeneous resident subsets, including Eomes^+^CD49a^+^ uterine trNK cells and CXCR6-associated populations which have been linked to tissue adaptation and pregnancy memory-like features^19,20^. However, classification based on tissue-residency marker expression remains imperfect in the decidua, because markers commonly associated with tissue residency in NK cells, such as CD49a, CD69, and CXCR6, may reflect retention or local adaptation without definitively distinguishing long-term resident cells from recently recruited and retained cells or from cells newly differentiated within the decidual niche. Thus, the unresolved question is not whether dNK cells traffic or reside, but rather how recruitment, local retention, and acquisition of tissue residency are temporally integrated across gestation.

Nevertheless, trafficking alone is unlikely to account for the full expansion of the dNK-cell compartment, indicating that additional local mechanisms contribute to dNK homeostasis. Across murine and human studies, evidence supports local proliferation as a contributor to dNK-cell expansion^5,21^. In mice, BrdU incorporation identified proliferating NK cells in the decidua, while Ki-67 and NKp46 co-staining provided histological evidence of proliferating NK cells within uterine tissue sections^5,22^. In humans, Immunohistochemical detection of Ki-67⁺CD56⁺ lymphocytes in first-trimester decidua provided further evidence of local NK-cell proliferation^23^. However, this expansion is not shared equally across NK subsets. Murine uterine trNK cells exhibited greater proliferative activity than cNK cells during decidualization^5^. Similarly, in human decidua, single-cell transcriptomics identified a proliferating NK-cell subset expressing MKI67 and TOP2A^24^, while mass cytometry identified a Ki-67-high dNK population^21^. Collectively, these findings support a model in which both trafficking and local proliferation contribute to dNK accumulation. However, the relative mechanistic contribution of proliferation versus recruitment remains unclear across gestation, particularly among trNK cells and migrated subsets.

A comparatively underexplored contributor to dNK-cell homeostasis is apoptosis. Human studies suggest that decidual lymphocytes in early pregnancy can engage FasL-dependent apoptotic pathways in addition to perforin-mediated killing^25,26^. In parallel, decidual stromal cells were shown to protect lymphocytes from apoptosis, indicating that local stromal cues may support dNK-cell survival by restraining cell loss^27^. Recently, Annexin V-based analyses linked altered miR-122-5p expression to changes in decidual NK-cell apoptosis in unexplained recurrent pregnancy loss^28^. However, most of this evidence addresses dNK-mediated apoptosis of target cells rather than apoptosis of dNK cells themselves, leaving the contribution of apoptosis to regulation of dNK homeostasis across gestation incompletely understood.

Another unresolved layer is the transcriptional program that links these cellular behaviors to dNK adaptation. Across murine and human species, T-bet (Tbx21) and Eomes (Eomesodermin) are the key regulators of NK-cell development and homeostasis^29,30^. In mice, combined loss of T-bet and Eomes impairs NK-cell development^31^, whereas in mature human peripheral NK cells, it disrupts NK-cell identity and induces an ILC3--like program^30^. Nevertheless, T-bet and Eomes have distinct but partially overlapping roles at different stages of NK-cell development^29,32^. In mice, Eomes promotes early lineage specification and immature NK-cell survival, whereas T-bet supports terminal maturation in murine conventional NK cells^29,32^. Human progenitor differentiation study supports a similar distinction between Eomes-associated early development and T-bet-associated terminal maturation^32^. In addition, T-bet regulates NK-cell trafficking by promoting S1PR5-dependent tissue exit from the bone marrow and lymph nodes^33^. However, T-bet and Eomes expression and functional requirements vary with tissue context. Human liver-resident NK cells are enriched for an Eomes-high, T-bet-low phenotype relative to their peripheral counterpart^34^. In virgin mice, uterine trNK cells can develop in the absence of T-bet^5^, whereas conditional deletion of Eomes in *Ncr1*-expressing cells eliminates uterine trNK cells in both virgin and pregnant mice, demonstrating an essential role for Eomes in uterine trNK-cell development^35^. Furthermore, T-bet deficiency increases the proliferative activity predominantly in mature murine NK cells, whereas immature CD11b⁻ cells are largely unaffected^29^. However, how T-bet expression relates to dNK-cell proliferation and tissue residency during pregnancy remains unclear. Our recent fate-mapping study demonstrated progressive and selective T-bet downregulation in dNK cells while maintaining Eomes protein expression. This adaptation was accompanied by reduced cytotoxic gene expression, IFN-γ-associated programs, and increased expression of adhesion-and homing-associated genes^36^. In this study, we hypothesized that dNK-cell residency is a dynamic state maintained through the coordinated contributions of trafficking, local proliferation, apoptosis, and remodeling of residency-associated phenotypes, while these processes are associated with T-bet expression in a gestational stage specific manner. Specifically, we asked:

a. Whether dNK cell residency-associated phenotypes and vascular accessibility change across gestation.
b. How trafficking, local proliferation, and apoptosis contribute to the maintenance and remodeling of the dNK-cell pool across pregnancy
c. How these cellular dynamics relate to selective and progressive downregulation of T-bet expression within dNK cells across pregnancy.

Elucidating this mechanistic distinction is critical, as aberrant dNK cell dynamics are implicated in defective placentation and a spectrum of adverse pregnancy outcomes^2,35^.

## 2. Materials and Methods

### 2.1. Mice

Female and male C57BL/6J (B6) mice were purchased from Jackson laboratory (Bar Harbor, ME, cat# 000664). Transgenic mouse model, loxP-flanked STOP-RFP reporter mouse strain (B6;129S6Gt(ROSA)26Sortm14(CAG-tdTomato)Hze/J; henceforth B6.Rosa, Jackson Lab, cat# 007914) were crossed with Tbet-Cre reporter strain (B6.CBA-Tg(Tbx21-cre)1Dlc/J, henceforth B6.Tbet^Cre^, Jackson Lab, cat# 024507) to produce B6.Rosa^RFP^/Tbet^Cre^ mouse model, respectively. These animals have a floxed STOP codon in front of the bright RFP transgene (tdTomato), which is removed (allowing RFP synthesis) in cells that have expressed the T-bet transcription factors at any prior time. B6.Tbet^Cre^ animals were backcrossed to C57Bl/6 for at least 2 generations at Jackson Laboratories and were backcrossed an additional 5 generations prior to breeding with B6.Rosa animals, with resulting Rosa/Tbet^Cre^ animals being backcrossed into the B6 background for at least 8 generations.

C57BL/6J (B6) and B6.Rosa^RFP^/Tbet^Cre^ Female mice (6-13 weeks) were used for timed mating and experiments. The day when a vaginal plug was detected in a timed mating was counted as gestational day 0.5. Virgin mice and the mice at various specified gestational day (early: 7, 8, mid:12, 13, 14, late: 16, 17,18 days) were sacrificed, and gestational day of decidua saved for analysis^36^. The number of mice used was summarized in Table 1.

**Table 1.**
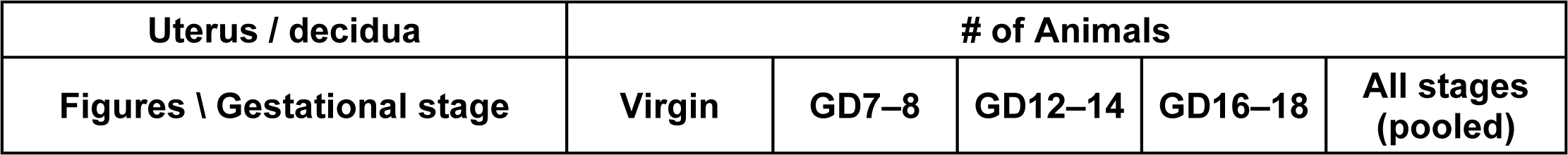
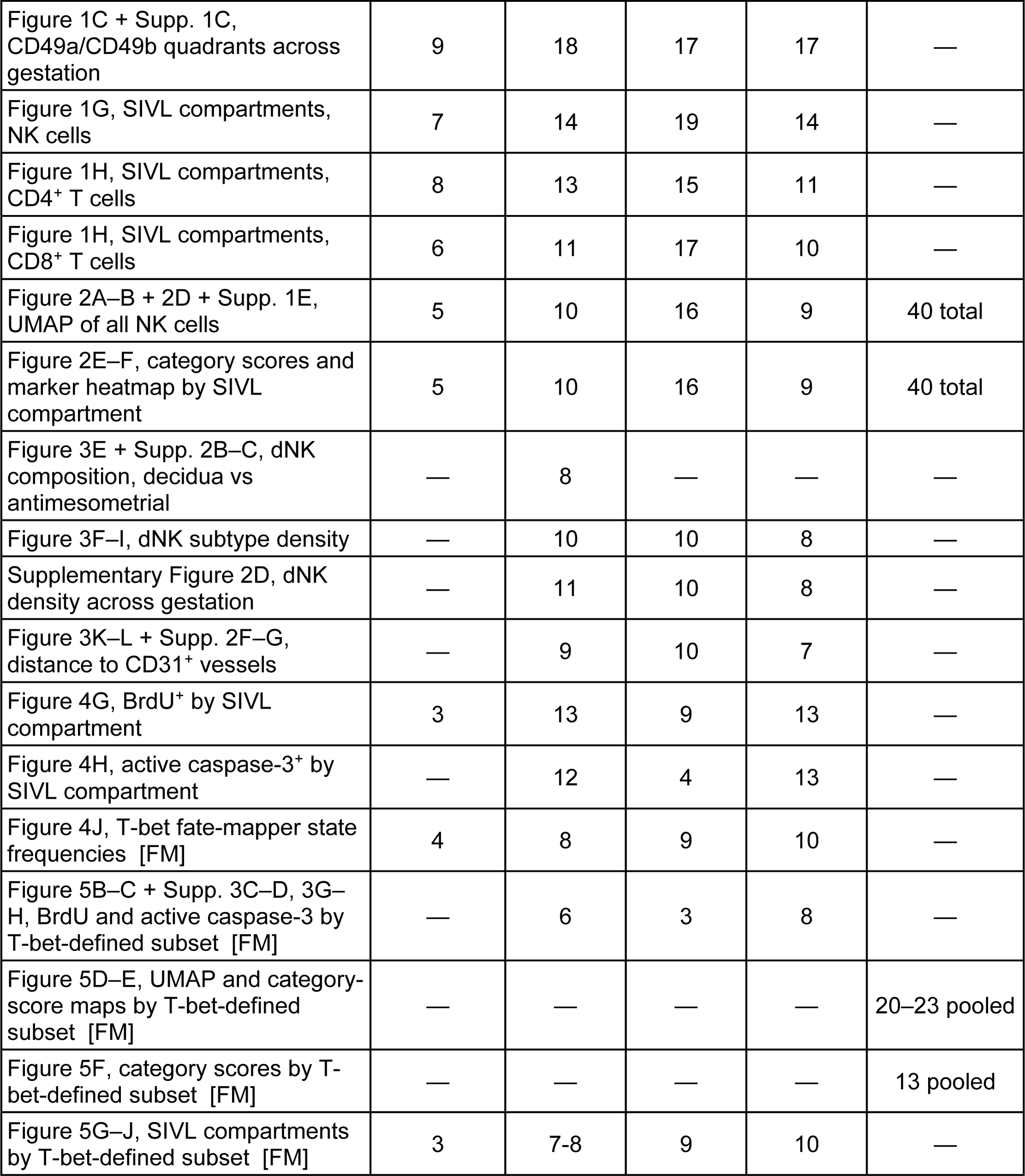
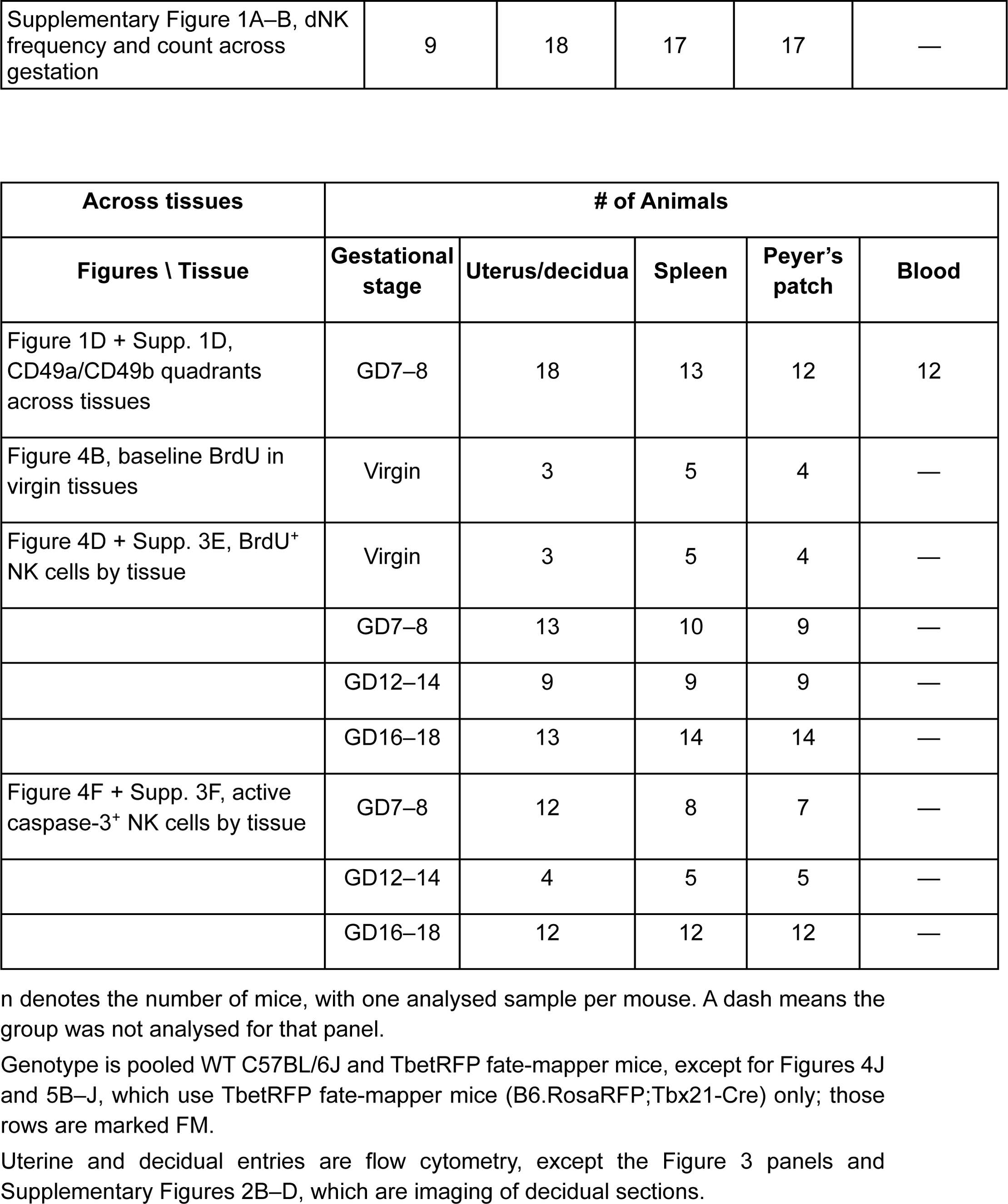
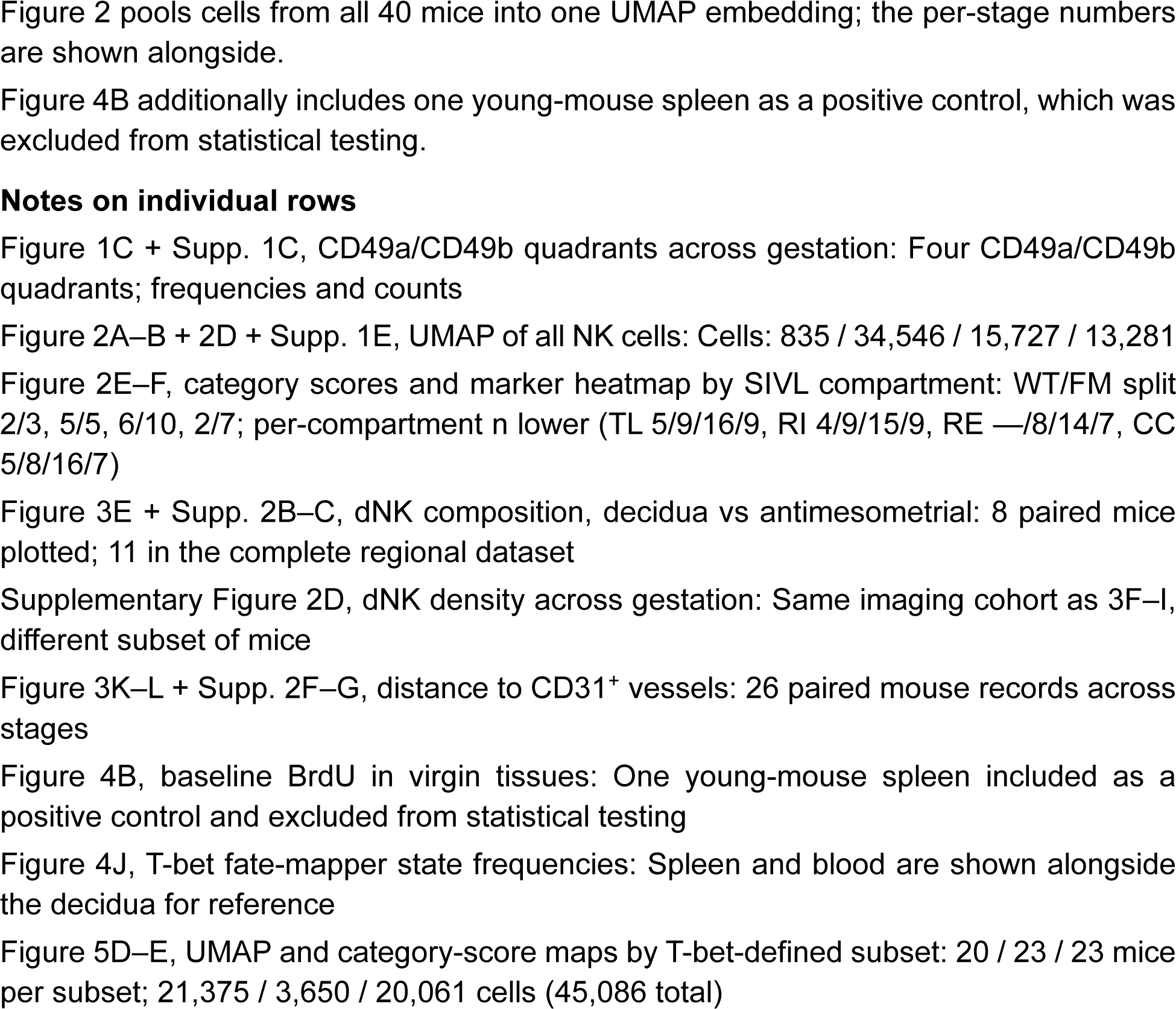
Number of animals for each figure panel.

### 2.2. Serial Intravascular Labeling (SIVL) and BrdU (5-bromo-2′-deoxyuridine) pulse-labeling of decidual leucocytes

Mice were intraperitoneally injected with 100 mg/kg body weight BrdU (eBioscience™ BrdU Staining Kit, cat# 8812-6600-42 for Flow Cytometry, BrdU, cat# 00-4440-51A) followed by 24 hours and 6.7 μg of AF488-conjugated CD45 antibody 24 hours and 6.7 μg of AF647-conjugated CD45 antibody 5 minutes before sacrifice. Virgin mice and pregnant mice were sacrificed, and the gestational day of each decidua was recorded^7^.

### 2.3. Tissue processing and immune cell isolation

Mouse deciduae were collected, minced with scissors in RMPI 1640 containing collagenase type V (Worthington Biochem, cat# LS005282)/DNAse I (Worthington Biochem, cat# LS006344). These specimens were then loaded in gentleMACS^TM^ C tube (Miltenyi Biotec Inc. San Diego, CA, cat# 120-005-331), and a specially adapted tissue dissociation program run in gentleMACS™ Dissociator for 12 minutes (Miltenyi Biotec Inc. San Diego, CA, cat# 130-096-427)^37^. Spleen and Peyer’s patches (used as control tissues) were mechanically dissociated in RMPI 1640 containing 10% heated FBS in gentleMACS^TM^ C tube, by running corresponding programs for different tissue types in gentleMACS™ Dissociator. After dissociation, homogenates were filtered through 70 μm cell strainer, and red cells of splenic specimens were lysed with ACK lysis buffer (Life Technologies, cat# A10492-01)^37^.

Peripheral blood was collected into EDTA-coated tubes and processed shortly after collection. Blood samples were diluted with sterile PBS and peripheral blood mononuclear cells (PBMCs) were isolated by density-gradient centrifugation using Ficoll-Paque (fisher scientific-Cat 45-001-751). Diluted blood was layered over Ficoll-Paque and centrifuged according to the manufacturer’s instructions without brake. The mononuclear cell layer at the plasma-Ficoll interface was collected and washed with PBS to remove platelets and residual Ficoll. Single cell suspension obtained was used for downstream flow-cytometric analysis.

### 2.4. Flow cytometry labeling

Single cell suspensions were first labeled with LIVE/DEAD® fixable blue stain (Invitrogen, cat# L34962) and subsequently a cocktail of flourochrome-conjugated monoclonal antibodies (list in Table 2) according to the manufacturer’s instructions. Briefly, antibodies were diluted in BD Horizon BrilliantTM Stain Buffer (BD Biosciences, San Jose, CA, cat# 566349) and used to label cells for 30 mins then washed. Cells were pulse-vortexed to ensure proper resuspension and fixed with BrdU fixation/permeabilization Buffer (eBioscience™, cat# 00-5223-56). The following day, cells were treated with DNase I working solution (eBioscience™, cat# 00-4425-10) according to the manufacturer’s instructions. After DNase treatment, cells were washed and stained intracellularly for transcription factors, including Eomes, T-bet and active caspase 3 followed by staining with fluorochrome-conjugated anti-BrdU antibody (Cat# 12-5071-42). Assessment for intracellular staining was done using BD Pharmigen™ Transcription Factor Buffer Set (BD, cat# 562574).

**Table 2.** Antibodies used for flow cytometry and multiplex immunofluorescence imaging.

| Panel |  |  |  |  |  |
|---|---|---|---|---|---|
| **Marker** |  | **Color/Format** | **Host/Target** | **Cat#** | **Clone** |
| CD161 |  | Brilliant Ultraviolet 395 | Mouse | 564144 | PK136 |
| CD19 |  | RealBlue 744 | Mouse | 570652 | 1D3 |
| CD49b |  | Brilliant Ultraviolet 661 | Mouse | 741523 | HMa2 |
| CD127 |  | Brilliant Ultraviolet 737 | Mouse | 612841 | SB/199 |
| CD3e |  | Pacific Blue | Mouse | 100334 | Bio145-2C11 |
| CD69 |  | Brilliant Violet 480 | Mouse | 746813 | H1.2F3 |
| CD103 |  | Brilliant Violet 711 | Mouse | 121435 | 2.00E+07 |
| CD45 |  | Alexa Fluor 488 | Mouse | 103122 | 30-F11 |
| CD49a |  | PerCP-Cy5.5 | Mouse | 564862 | HA31/8 |
| CD4 |  | PE-Fire 700 | Mouse | 100483 | GK1.5 |
| CD45 |  | Alexa Fluor 647 | Mouse | 103124 | 30-F11 |
| Ly-6G |  | Alexa Fluor 700 | Mouse | 557979 | RB6-8C5 |
| CD45 |  | APC-Fire 810 | Mouse | 103173 | 30-F11 |
| CD11c |  | Alexa Fluor 700 | Mouse | 560583 | HL3 |
| EOMES |  | PE-Cy7 | Mouse | 25-4875-80 | Dan11mag |
| T-bet |  | Brilliant Violet 421 | Mouse | 563318 | O4-46 |
| CD8a |  | RealYellow 586 | Mouse | 568164 | 53.67 |
| CD62L |  | NovaFluor Blue 610 / 30S | Mouse | M006T02B05-A | MEL-14 |
| CD184 (CXCR4) |  | Brilliant Violet 650 | Mouse | CBMAB-C0826-LY | 2B11/CXCR4 |
| S1PR5 |  | Alexa Fluor 405 | Mouse | FAB9084MFV610 | 1196A |
| BrdU |  | PE | Mouse | 12-5071-42 | BU20A |
| Live/Dead Fix Blue |  | Live/Dead Fix Blue | All Species | L34962 |  |
| Caspase 3 (active) |  | RealBlue 780 | Mouse | 570335 | C92-605.rMAb |
| CCR6 |  | Brilliant Violet 785 | Mouse | 129823 | 29-2L17 |
| **antibody table for multiplex immunofluorescence** |  |  |  |  |  |
| **Target / marker** |  | **Purpose in panel** | **Fluorophore** | **Clone** | **Host / isotype** |
| CD45 |  | Leukocyte labeling for SIVL: 24-h in vivo label | Alexa Fluor 488 | 30-F11 | Rat IgG2b, κ |
| CD45 |  | Leukocyte labeling for SIVL: 5-min in vivo vascular label | Alexa Fluor 647 | 30-F11 | Rat; monoclonal; vendor lists concentration 0.5 mg/mL |
| CD31 / PECAM-1 |  | Endothelial/vessel marker | Alexa Fluor 555 | D8V9E | Rabbit IgG; monoclonal |
| NCR1 / NKp46 |  | NK-cell marker | Alexa Fluor 350 | Polyclonal | Goat IgG; polyclonal |

### 2.5. Data Analysis

Spectral flow cytometry data were acquired using Cytek® Aurora spectral cytometer. Cytek Aurora is a five-laser, 67-channel spectral flow cytometer configured as follows: 355 nm (16 channels), 405 nm (16 channels), 488 nm (14 channels), 561 nm (10 channels), and 640 nm (8 channels). Spectral data were unmixed using SpectroFlo® software (version 3.2.1) with a reference matrix generated from single-stained controls.

Manual gating analysis was performed using FlowJo v10.10 (BD Biosciences, Franklin Lakes, NJ, USA). Statistical analysis was conducted manually using GraphPad Prism version 7 (GraphPad Software Inc., La Jolla, CA, USA) as well as R version (4.5.1 (2025-06-13 ucrt) in RStudio version (2023.12.1.402). Statistical significance among SIVL-defined dNK cells was assessed using a linear mixed-effects model followed by Tukey-adjusted post hoc pairwise comparisons (p < 0.05). All the statistical details were summarized in Table S1.

### Calculation of composite trafficking-associated programs scores

Composite marker scores were calculated to summarize trafficking-and tissue residency-associated programs in dNK cells. Three marker categories were defined a priori: tissue retention/residency program, comprising CD49a, CD69, and CD103; circulatory/egress-like, comprising CD49b, CD62L, and S1PR5; and chemokine responsiveness program, comprising CXCR4 and CCR6. Using the transformed expression matrix applied for UMAP and heatmap analyses, each marker value was independently standardized across the complete dNK-cell dataset to a mean of zero and a standard deviation of one. For each cell, the tissue-retention/residency score was calculated as the arithmetic mean of the standardized CD49a, CD69, and CD103 expression values; the circulatory/egress-like score was calculated as the mean of standardized CD49b, CD62L, and S1PR5 expression; and the chemokine-response score was calculated as the mean of standardized CXCR4 and CCR6 expression. Thus, each dNK cell received one score for each marker category, with all markers contributing equally.

For graphical summaries across gestational stages and SIVL compartments, cell-level category scores were averaged within each mouse and SIVL compartment, and group means and standard errors were calculated from these mouse-level means. For comparisons among SIVL compartments, separate linear mixed-effects models were fitted for each category score, with SIVL compartment included as a fixed effect and mouse included as a random intercept to account for repeated compartment measurements within the same animal. Pairwise contrasts were obtained from estimated marginal means and adjusted for multiple comparisons using the Holm method. The same approach was used to compare T-bet-defined states, where T-bet-defined state included as the fixed effect and mouse as the random intercept. Individual mice, rather than cells, were treated as independent biological replicates.

### Spatial mapping of SIVL-defined-dNK cells by multiplex immunofluorescence

Spatiotemporal mapping of dNK cells was assessed using multiplex immunofluorescence staining of paraffin-embedded tissue sections. Experimental and control slides were processed in parallel under identical staining conditions. Briefly, tissue biopsies were collected and fixed overnight in 4% paraformaldehyde and paraffin-embedded (FFPE). FFPE tissue sections (5 μm) were first deparaffinized through 2 sequential xylene washes (Fisher Chemical, Ref X3P-1GAL, Lot 207995) for 5 minutes each, followed by graded ethanol rehydration (Fisher Chemical by Decon Labs Inc., cat#190, proof 2801, 200 proof 2701) for 5 minutes each. Antigen retrieval was performed using 10 mM sodium citrate buffer, pH 6.0. (Novus Biologicals. Ref NB900-62075, Lot X068-M-A.) for 10 minutes. Sections were then washed and permeabilized using 1% FBS in 0.4% PBS-Triton, followed by blocking with 5% FBS in 0.4% PBS-Triton (Fisher BioReagents.cat# BP337-100, Lot 163454) at room temperature for 30 minutes. Serial sections were stained with either hematoxylin and eosin and multiplex immunofluorescence staining. For multiplex immunofluorescence staining, directly conjugated primary antibodies were diluted in 1% FBS/0.4% PBS-Triton and applied to tissue sections (list in Table 2). The antibody cocktail solution was added, and slides were incubated for 1-2 hours at room temperature and stored overnight flat in a humidified chamber at 4°C then washed twice with 1% FBS in 0.4% PBS-Triton 10 minutes each. Nuclear counterstaining was performed using NucSpot 750/780, 1000x in DMSO: Biotium, Cat# 41038-T), and sections were incubated for 10 minutes then mounted using antifade mounting medium (EverBrite Hardset Mounting Medium, Biotium, Cat# 23003).

### Image acquisition and regions

Histology sections were imaged on a Leica’s THUNDER Imager 3D Assay using a HC PL APO 40x/0.95 CORR objective (resolution of 0.327 µm) using the large image and automatic stitching options.

Tail vein CD45 injections at 24 h were labeled with Alexa Fluor 488 and detected using the TEAL511/B535 channel, whereas CD45 injections after 5 min were labeled with Alexa Fluor 647 and detected using the RED635/D642 channel. These fluorophore-channel combinations provided robust signals and clear spectral separation, which was central to this study. NCR1/NKp46 conjugated to Alexa Fluor 350 was detected using the VIOLET395/A460 channel, and CD31 conjugated to Alexa Fluor 555 was detected using the YELLOW575/C590 channel to visualize the vasculature. The nuclear stain was detected using the NIR730/757-780 channel to minimize spectral overlap and channel crosstalk with the other fluorophores, particularly given the abundance of the nuclear signal. Fluorophore-target assignments and detection channels were optimized to enable unambiguous identification of the labeled structures (more details in upcoming subsections), and signal specificity and channel separation were experimentally validated by iterative staining of control slides alone and in combination.

All quantifications were performed in QuPath 0.6.0 using custom Groovy scripts, which are included with the supplementary method. Images acquired at lower magnification were excluded, because at those pixel sizes the spot-smoothing step of the particle filter falls below half a pixel and is disabled, so the filter applied is not the same filter.

Two implantation regions were annotated manually per section, the decidua and the antimesometrium, and each was measured as a single whole region. Sample and control regions were drawn by the same criterion.

### Cell segmentation and NCR1 gating

Nuclei were segmented with StarDist (dsb2018_heavy_augment model, probability threshold 0.50, 0.25 µm working resolution) within both annotated regions. Each nucleus was expanded isotropically by 3.0 µm to approximate the cell body, and nuclei exceeding 70 µm² were discarded as segmentation merges. A cell was considered as an eligible candidate (dNK cell) if its mean NCR1 intensity exceeded the region mean plus two standard deviations, a gate computed independently for each region so that section-to-section differences in staining intensity do not alter the criterion. Positivity is expressed throughout as a percentage of eligible dNK cells.

### Tracer signal detection and threshold setting (anti-CD45 AF488 24h injection and antiCD45 AF647 5 min Injection)

Tracer signals were detected as localized regions of increased fluorescence relative to the surrounding tissue, using a difference-of-Gaussians filter. Each tracer channel was smoothed with a Gaussian filter (σ = 0.25 µm), and a local background estimate was obtained by smoothing the same channel with a broader Gaussian (σ = 3.0 µm). The background was subtracted from the smoothed image, and the resulting image was retained as a signed residual, so pixels below their local background carry negative values. Pixels above threshold formed a mask, touching regions were segmented by a Euclidean-distance watershed, and the resulting individual regions were labelled using 8-connectivity. Regions between 0.30 and 8.00 µm² were retained. The upper limit excludes erythrocytes, which are autofluorescent in the far red and form far larger regions. In a vascular-rich uninjected control, these regions reached 143.85 µm², exceeding the size of any candidate region detected in an injected section.

### Thresholds

For channel AF488, the detection threshold was set to eight times each section’s noise scale (8σ). Noise scale (σ) was estimated within each image tile as a robust standard deviation, calculated as 1.4826 × the median absolute deviation (MAD) of the filtered residual. The section-level noise scale was then defined as the 25th percentile of these tile-level estimates. This lower quantile (25^th^ percentile) was used because bright objects increased within-tile variability, and a pooled estimate overestimated σ ∼ 2-fold in object-rich sections and disproportionately raising thresholds in sections with stronger tracer signal. Thresholds were therefore scaled to each section’s background variability. Across four uninjected decidual and antimesometrial controls, 0.78–1.12% of background pixels exceeded the threshold, although the absolute thresholds ranged from 274 to 1,324.

For channel AF647, a fixed reference was derived from uninjected controls (decidua and animesometrial region) by quantile matching (170.8 decidua, 155.0 antimesometrium), and the threshold was the greater of that reference and 25 times the section’s own noise scale (25σ), with σ estimated as described above. This value was compared with a fixed reference derived by quantile matching: 170.8 for the decidua and 155.0 for the antimesometrial region. The higher of the two values was used as the final detection threshold.

### Spectral specificity filtering

Tissue autofluorescence is broadband and raises all channels together, while tracer signal is confined to its own channel. Each eligible candidate object in both tracer channels was therefore tested for channel specificity.

For each candidate signal region, fluorescence in each channel normalized as (S = peak/ σ) (Eq. 1), where peak was the maximum intensity within that region in the corresponding channel. A dominance score was then calculated as (D = min (S_CD31, S_NCR1) / S_tracer) and regions D ≤ 0.50 were retained. This filter was applied because the tissue autofluorescence is broadband and raises every channel together, which can increase signal across channels and yield D values near 1 when the normalized intensities are comparable.

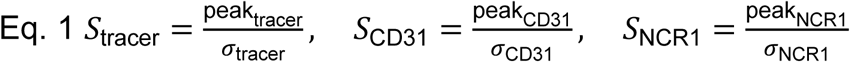

In the AF488 channel, the filter rejected 100% of candidate objects in all four uninjected controls (at a median D of 0.797–0.896, and 8.5–17.7% of candidates in injected samples, where retained objects had a median D of 0.032–0.283). Because AF647 emission lies outside the principal autofluorescence band, the filter rejects little there (0.5% of candidates on a representative section, retained median D = 0.016).

Then a cell was classified as positive for a tracer channel if it contained at least one region of increased fluorescence intensity retained after autofluorescence filtering (D ≤ 0.50) and no more 35% of pixels within its nuclear mask exceeded the channel-specific intensity threshold. The second exclusion criterion is cells with extensive above-threshold tracer-channel fluorescence overlapping the nuclear mask.

### Controls, exclusions and validation

Four uninjected, unstained two decidual and two antimesometrial control sections, spanning a 4.8-fold range of AF488 noise scale, were processed identically to the samples. All four returned 0.000 % AF488-positive cells and zero retained AF488 particles.

Before quantification, every channel of every section was profiled in a single read-only pass across the whole project, and all sections were compared side by side. A channel carries considerable signal only when its bright tail stands clear of its own background. Specifically, in a live channel the 99.9th percentile lies well above the median, whereas in a dead channel the median, the 99.9th percentile and the maximum all together fall close to each other and close to zero, so sections in which a tracer channel (AF488 or AF647) or the NCR1 channel showed no such separation were excluded. This is a channel without dynamic range and cannot be measured at any threshold. These exclusions are properties of the image alone: the profiling pass computes percentiles only, never positivity, and was completed before any threshold was applied. Accordingly, the analysis rejects regions with flat fluorescence profiles, defined as those with variance equals. To further confirm that the reported values do not depend on where a region was drawn, one control section was measured under four combinations of threshold method and region: under a fixed threshold its value varied six-fold with region placement (5.366 % and 0.905 %), whereas under the reported method nearly doubling the region area changed nothing (0.000 % in both cases).

### Comparison of SIVL-dNK-cell enrichment across gestation

As a complementary analysis, tracer positivity was assessed using a relative threshold set at the 99.5th percentile of per-cell tracer intensities within each image and region, followed by the cell-level dominance filter (D ≤ 0.50). NCR1 served as a reference channel in this filter without changing the threshold used to identify dNK cells (median intensity+2SD). This combination was selected based on uninjected control sections and retained positive cells in injected samples. This approach was used because absolute thresholds varied from 146 to 1,324 across sections. DNK cells were identified using the established NCR1 positivity threshold of the median plus two standard deviations.

### Experimental unit and sample sizes

Serial sections from the same sample were summed within each mouse before statistical testing, and all reported statistics were calculated using per-mouse values. Cells within a section were not considered independent biological observations because they shared the same mouse, vascular bed, region annotation, and segmentation procedure.

Due to small sample size, nonparametric tests were selected. Gestational SIVL-defined-dNK subset comparisons were performed using the Kruskal-Wallis test followed by pairwise Wilcoxon rank-sum tests with Holm correction across the three pairwise comparisons

### Distance measurement to nearest CD31^+^ blood vessel. Vessel segmentation

CD31-positive vessels were segmented using image-specific Otsu thresholding following background subtraction. Diffuse background fluorescence was removed by subtracting a Gaussian-blurred copy of the CD31 image (σ = 50 µm) from the original image. An intensity threshold was then calculated independently from each image’s background-corrected CD31 intensity distribution using Otsu’s method image. Because CD31 background intensity varied between sections, image-specific thresholds accommodated these variations. Vessel contours were extracted at this threshold and connected components smaller than 20 µm² were removed as small noise components. The resulting segmentation was clipped to the decidual region. To avoid the edge artifact, the region was eroded inward by 40 µm to exclude CD31 signal within 40 µm of its boundary before clipping. These measures were added to the Qupath custom scripts to use the segmented CD31-positive vessel-wall signal as the reference for distance measurements, rather than a filled vessel body.

### Distance measurement

For every detected nucleus, the distance to the nearest CD31 annotation was computed with QuPath’s detection. Specifically, the distance was measured from the nucleus centroid to the nearest boundary of the nearest CD31^+^ annotation. Distance is calculated from the vascular staining centroid.

### To analyze RI enrichment at distance 0 (vessel contact)

Decidual per-cell vessel distances were exported in two datasets: one is restricted to SIVL-defined-dNK cells classified as RI, RE, CC, or TL, and another containing every detected nucleus. The latter provided the non-dNK reference population required for RI enrichment analyses at distance 0. Region areas and total nuclei per region were exported separately to calculate cell density (cells/mm²) and cell abundance normalized to 10,000 nuclei.

### Statistical analysis

AF488 and AF647 Tracer signals was quantified using two complementary endpoints: the percentage of eligible cells classified as tracer-positive and the density of retained tracer particles per mm². Differences in the frequency of tracer detection between experimental and control groups were assessed using Fisher’s exact test. Quantitative sample-versus-control differences were assessed using the same rank test. All statical analysis was provided in Table S1

## 3. Results

### 3.1. Pregnancy alters CD49a/CD49b-defined dNK phenotypes

We first asked whether pregnancy alters classical CD49a/CD49b-defined uterine NK-cell phenotypes. We analyzed CD45+ immune cells from nonpregnant virgin uterus and pregnant decidua of C57BL/6J (B6) and B6.RosaRFP/TbetCre fate mapper mice at early, mid, and late gestation (GD7-8, GD12-14, and GD16-18). Gestational changes were determined in the overall dNK-cell compartment by quantifying both dNK-cell frequency within the CD45+Lin^neg^ immune compartment and total dNK-cell abundance across pregnancy. Compared with the virgin uterus, pregnant decidua showed a marked accumulation of dNK cells, particularly at early gestation (Supplementary Fig.S1A and B). We next examined whether pregnancy-associated dNK accumulation was accompanied by changes in CD49a/CD49b-defined NK cell phenotypes. Viable CD45⁺Lin^neg^NK1.1⁺ cells were identified as dNK cells and further stratified by CD49a and CD49b expression to distinguish tissue resident-like CD49a⁺CD49b⁻ dNK cells, conventional-like CD49a⁻CD49b⁺ NK cells, and mixed/transitional double-positive CD49a⁺CD49b⁺ dNK cells (Fig. 1A). CD45+Lin^neg^NK1.1+ cells revealed marked remodeling of CD49a/CD49b-defined subsets, particularly the early gestational enrichment of CD49a⁺CD49b⁺ dNK cells (Fig. 1B, C and supplementary Fig.S1C). Tissue comparison at GD7-8 showed that this phenotype was most prominent in the decidua compared with peripheral tissues supporting a tissue-specific decidual remodeling program (Fig. 1D and Supplementary Fig.S1D).

**Figure 1.**
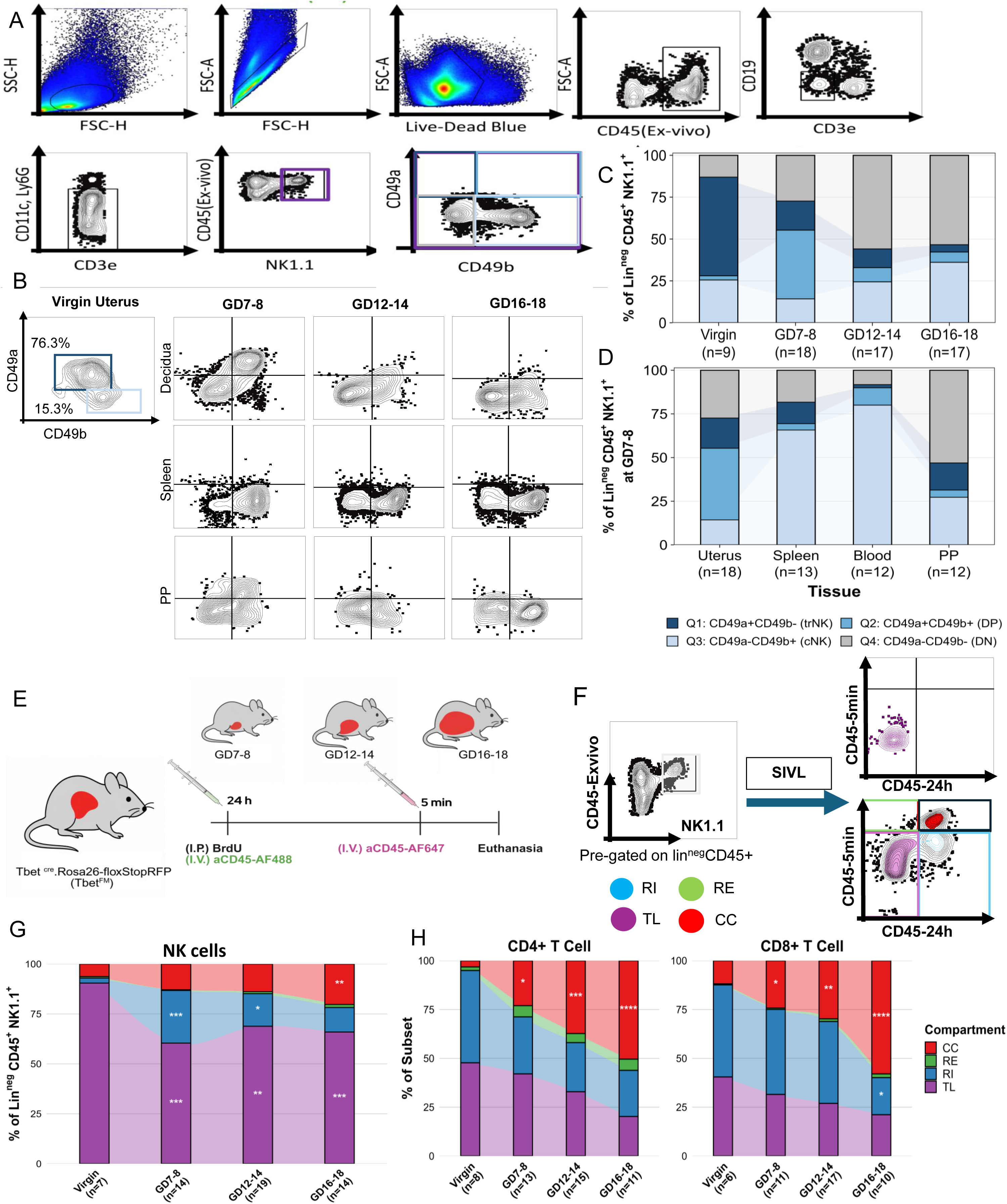
Pregnancy remodels uterine/decidual CD49b/CD49a NK-cell subset composition and trafficking dynamics. **(A)** Representative flow-cytometric gating strategy to identify live singlet CD45⁺Lin⁻NK1.1⁺ uterine/decidual NK cells and resolve CD49a/CD49b-defined populations. **(B)** Representative CD49a-versus-CD49b profiles of NK cells from the virgin uterus and from decidua, spleen, and Peyer’s patches (PP) across pregnancy. Numbers indicate the percentage of cells within the indicated parent population. **(C)** Mean composition of CD49a⁺CD49b⁻, double-positive CD49a⁺CD49b⁺, CD49a⁻CD49b⁺, and CD49a⁻CD49b⁻ NK-cell populations in the virgin uterus and decidua across gestation (virgin, *n* = 9; gestational days (GD) 7-8, *n* = 18; GD12-14, *n* = 17; GD16-18, *n* = 17). **(D)** Mean CD49a/CD49b-defined NK-cell composition in the uterus/decidua, spleen, blood, and PP at early pregnancy GD7-8 (uterus, *n* = 18; spleen, *n* = 13; blood, *n* = 12; PP, *n* = 12). Statistical analysis: Kruskal-Wallis. Pairwise test: Dunn post hoc with Benjamini-Hochberg false-discovery-rate adjustment. Significance: ns, p≥0.05; *, p<0.05; **, p<0.01; ***, p<0.001; ****, p<0.0001. **(E)** Experimental design. T-bet fate-mapping mice (Rosa/Tbet^Cre^mice) and C57BL/6J were analyzed at virgin and GD 7-8, 12-14, and 16-18. intraperitoneal (I.P.) BrdU and tail vein injection (I.V.) anti-CD45-AF488 were administered 24 hours before tissue collection, followed by (I.V) anti-CD45-AF647 5 minutes before euthanasia. **(F)** Representative gating strategy for serial intravascular labeling (SIVL) of NK cells. Based on 24-h and 5-min intravascular anti-CD45 labeling, cells were classified as tissue localized (TL; CD45-24 h⁻CD45-5 min⁻), recently intravascular (RI: CD45-24 h⁺CD45-5 min⁻), recent emigrant (RE: CD45-24 h⁻CD45-5 min⁺), or continuously circulating (CC: CD45-24 h⁺CD45-5 min⁺). **(G)** Mean distribution of SIVL-defined TL, RI, RE, and CC compartments among uterine/decidual NK cells at virgin and across pregnancy. **(H)** Corresponding gestational distributions of SIVL-defined compartments among uterine/decidual CD4⁺ and CD8⁺ T cells. Stacked bars represent group mean proportions normalized to 100%. Statistical analysis used a linear mixed-effects model with experiment as a random intercept, followed by Holm-adjusted pairwise comparisons of estimated marginal means. Stars: each compartment versus Virgin. * p<0.05, ** p<0.01, *** p<0.001.

These findings suggest that CD49a/CD49b-defined tissue-resident and conventional dNK-cell phenotypes are not fixed states in the decidua, yet they are actively remodeled across gestation. How the dNK remodeling is mediated is not clear, so we next asked whether it reflects changes in vascular accessibility, and trafficking-associated receptors and ligands programs.

### 3.2. Serial intravascular labeling (SIVL) of CD45 reveals distinct dNK-cell trafficking patterns accompanying pregnancy-associated remodeling

To determine how NK trafficking contribute to the remodeling observed, we applied SIVL approach we recently adapted to decidual study to classify CD45+Lin^neg^NK1.1⁺ NK cells and CD45⁺CD3⁺ T-cell subsets into four vascular-accessibility compartments: unlabeled tissue-localized cells (TL), cells labeled by the antibody injected 24-hour prior to but not the 5 minutes prior to sacrifice consistent with recent tissue entry (RI), cells labeled only by the 5-minute antibody consistent with recent vascular exposure (RE), and cells labeled by both antibodies indicating continuous vascular accessibility (CC) (Fig.1E and F)^7,38^. Quantification of the SIVL compartments revealed that, in the virgin uterus, NK cell pool was dominated by TL cells, whereas early pregnancy showed a clear expansion of the RI NK cells. As pregnancy progressed, TL dNK cells remained the major NK-cell compartment, but the relative contribution of circulation-accessible NK cells, particularly CC compartment, increased toward late gestation (Fig.1G). In contrast, CD4^+^ and CD8^+^ T cells showed a stronger progressive shift toward circulation-accessible compartments, with a marked late-gestation increase in CC cells, indicating the distinct trafficking patterns of lymphocytes as gestation advances (Fig.1H).

### 3.3 SIVL-defined dNK compartments exhibit distinct trafficking-and residency-associated programs that evolve across pregnancy

To further determine if trafficking alters the biological phenotype of dNK cell across gestation, we next asked whether SIVL-defined compartments carried distinct biological programs. To address this question, trafficking- and residency-associated markers were grouped into three categories: tissue-retention/residency-associated CD49a, CD69, and CD103; circulatory/egress CD49b, CD62L, and S1P5R; and chemokine-response CXCR4 and CCR6. For each cell, relative category scores were calculated for tissue retention/residency, a circulatory/egress-like phenotype, and chemokine responsiveness. The gated NK cells from virgin, GD7-8, GD12-14, and GD16-18 uteri/deciduae were jointly projected by UMAP using the phenotypic markers including CD49a, CD49b, CD103, CD69, CXCR4, CCR6, S1PR5 and CD62L. The same UMAP coordinates were then annotated independently by the gestational stage and SIVL compartments (Fig. 2A and B).

**Figure 2.**
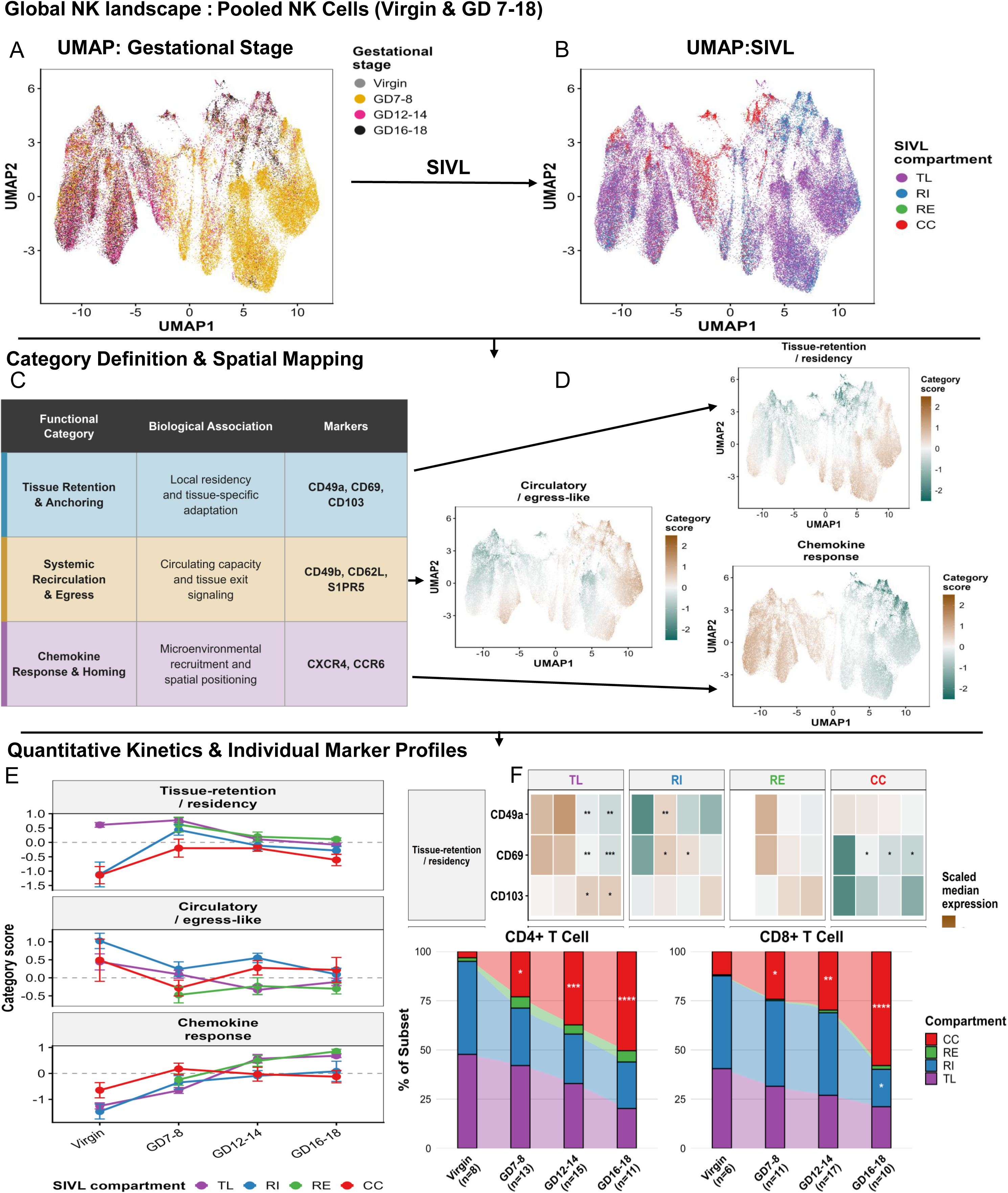
SIVL-defined dNK-cell compartments exhibit distinct trafficking-associated phenotypes across gestation. **(A)** representative UMAP of concatenated flow cytometry data of uterine/decidual NK cells across virgin and gestational days (GD 7-8, GD 12-14, and GD 16-18) based on the expression of the trafficking associated molecules: retention/residency (CD49a, CD69, and CD103), circulatory/egress-like phenotype (CD49b, CD62L, and S1PR5), and chemokine responsiveness (CXCR4 and CCR6). UMAP projection colored by gestational age: virgin, GD7-8, GD12-14, or GD16-18. **(B)** The same UMAP colored according to SIVL-defined compartment: TL, RI, RE, and CC. **(C)** Trafficking-associated markers were organized into three functional categories: tissue retention/residency (CD49a, CD69, and CD103), circulatory/egress-like phenotype (CD49b, CD62L, and S1PR5), and chemokine responsiveness (CXCR4 and CCR6). The categories respectively represent marker profiles associated with local retention, circulating NK-cell and egress potential, and responsiveness to chemokine signals **(D)** Distribution of the tissue-retention/residency, circulatory/egress-like, and chemokine-response category scores across the UMAP. For each cell, individual marker-expression values were standardized and averaged within their assigned category; brown and green indicate the relative higher and lower category scores, respectively. **(E)** Category-score trajectory: Mean tissue-retention/residency, circulatory/egress-like, and chemokine-response scores for each SIVL-defined compartment across reproductive stages. Data are presented as mean ± SEM. Colors denote TL (purple), RI (blue), RE (green), and CC (red). **(F)** Heatmaps showing scaled median expression of each constituent marker across reproductive stages within TL, RI, RE, and CC compartments. Brown and green indicate the relative expression above and below the marker-centered mean, respectively. Statistical analysis used a linear mixed model with estimated marginal mean contrasts against the virgin group and Holm adjustment for multiple comparisons. Data are presented as mean ± standard error of the mean. Asterisks in panels E and F indicate significant differences relative to the virgin group within the corresponding SIVL compartment: *P* < 0.05, **P** < 0.01, and ***P*** < 0.001. The number of mice per group is provided in Table 1.

Mapping by gestational stage showed extensive overlap among groups, indicating that dNK cells occupy a shared phenotypic continuum across pregnancy. Nevertheless, gestational stage-associated regional biases were evident. GD7-8 dNK cells were concentrated within the large right-hand region of the UMAP, whereas GD12-14 dNK cells were enriched across the central bridging region. In contrast, GD16-18 dNK cells were more prominently represented in the left and upper regions. Virgin-derived NK cells were visually sparse and did not form an isolated phenotypic region. Thus, pregnancy progression was associated with a redistribution of dNK cells across shared phenotypic states rather than the emergence of gestational stage-restricted populations.

Interestingly, SIVL compartments were also distributed across the entire UMAP. TL and RI dNK cells in particular were found in all major UMAP regions and exhibited extensive phenotypic overlap. By comparison, CC cells were predominantly co-clustered, whereas RE dNK cells were few and thus, visually sparse.

Projection of category scores onto the shared UMAP revealed continuous spatial gradients rather than discrete populations defined by a single program (Fig. 2C, D and individual marker-expression UMAPs; supplementary Fig.S1E).

Regions enriched for chemokine responsiveness showed higher CXCR4/CCR6 scores but lower tissue-retention and circulatory/egress scores, consistent with greater potential for chemokine-guided recruitment and positioning. By contrast, tissue-retention and circulation/egress scores partially overlapped, suggesting that tissue-adaptation and circulatory-associated features can coexist within the same phenotypic space, while intermediate regions exhibited mixed profiles. Thus, SIVL-defined compartments encompass heterogeneous and overlapping phenotypic states characterized by different combinations of tissue-association, circulatory/egress-related, and chemokine-responsiveness programs.

We next asked whether marker-defined tissue-retention, circulatory/egress-like, and chemokine-receptor phenotypes evolve across the pregnancy within each SIVL compartment. The tissue-retention/residency score demonstrated distinct compartment- and stage-associated trajectories (Fig. 2E and F). TL dNK cells maintained a positive retention/residency score in the virgin uterus and at GD7-8, followed by a reduction at mid-to-late gestation. RI and RE cells shifted from low scores in the virgin uterus to transient positive values at GD7-8. In contrast, CC compartment maintained a comparatively low tissue retention potential characterized by low CD69 and CD103 expression throughout pregnancy. The early retention-associated phenotype of TL cells and the transient increase observed in RI and RE cells was primarily associated with higher CD49a and CD69 expression, whereas CD103 expression was higher at mid and late gestation. Thus, the reduction in the retention/residency score did not reflect uniform loss of all residency-associated features but instead indicated a gestational shift from a CD49a/CD69-dominant phenotype toward greater CD103 representation.

The circulatory/egress-like score also varied by SIVL compartment as well as the gestational stage. RI dNK cells exhibited the highest overall scores, particularly at mid-gestation, corresponding to relatively high CD49b and CD62L expression. However, S1PR5 expression within RI cells progressively declined across gestation. TL compartment displayed a reduction in the circulatory/egress-like score after the virgin stage, predominantly associated with lower CD49b and S1PR5 expression, while CD62L was comparatively maintained. RE dNK cells retained a relatively low composite score despite increased S1PR5 expression at mid and late gestation. CC dNK cells showed a distinct pattern in which CD49b increased at mid and late gestation, whereas CD62L and S1PR5 remained low.

The chemokine-receptor expression score revealed a progressive gestational increase in TL and RE compartments. This transition was associated with increased CXCR4 and, more prominently, CCR6 expression at mid and late pregnancy. RI uterine/decidual NK subset shifted from a markedly low chemokine-receptor score in the virgin uterus toward values near the reference level during pregnancy. In contrast, CC dNK cells exhibited a modest, transient increase at GD7-8 without a sustained increase at mid or late gestation. Collectively, these results demonstrate that SIVL compartments are phenotypically dynamic and undergo distinct gestational remodeling of tissue-association, circulatory/egress-like, and chemokine-receptor profiles across pregnancy. Therefore, we next asked whether distinct phenotypic programs among SIVL-defined dNK populations were reflected in their spatial distribution within the decidua or in their proximity to the decidual vasculature.

### 3.3. Spatial imaging validates persistent perivascular enrichment of SIVL-defined RI dNK across gestation

To define anatomically the dNK vascular accessibility and trafficking contribution to tissue compartments we performed multiplex immunofluorescence on uterine tissue sections from GD7-8, GD12-14 and GD16-18 pregnancies (Fig.3A and supplementary Fig.S1F-G,). In decidual tissue sections from SIVL-injected mice, NCR1 staining identified dNK cells, while CD45 labeling distinguished SIVL-defined compartments (CC, RI, TL, RE) and was undetectable in controls (Fig. 3B, C and supplementary Fig. S2A). At GD7-8, Fluorescence-based anatomical segmentation, guided by serial H&E sections, was used to define mesometrial decidua basalis and antimesometrial decidua capsularis (Fig. 3D) and subsequently anatomically partition the trafficking contribution seen on aggregate flow studies.

**Figure 3.**
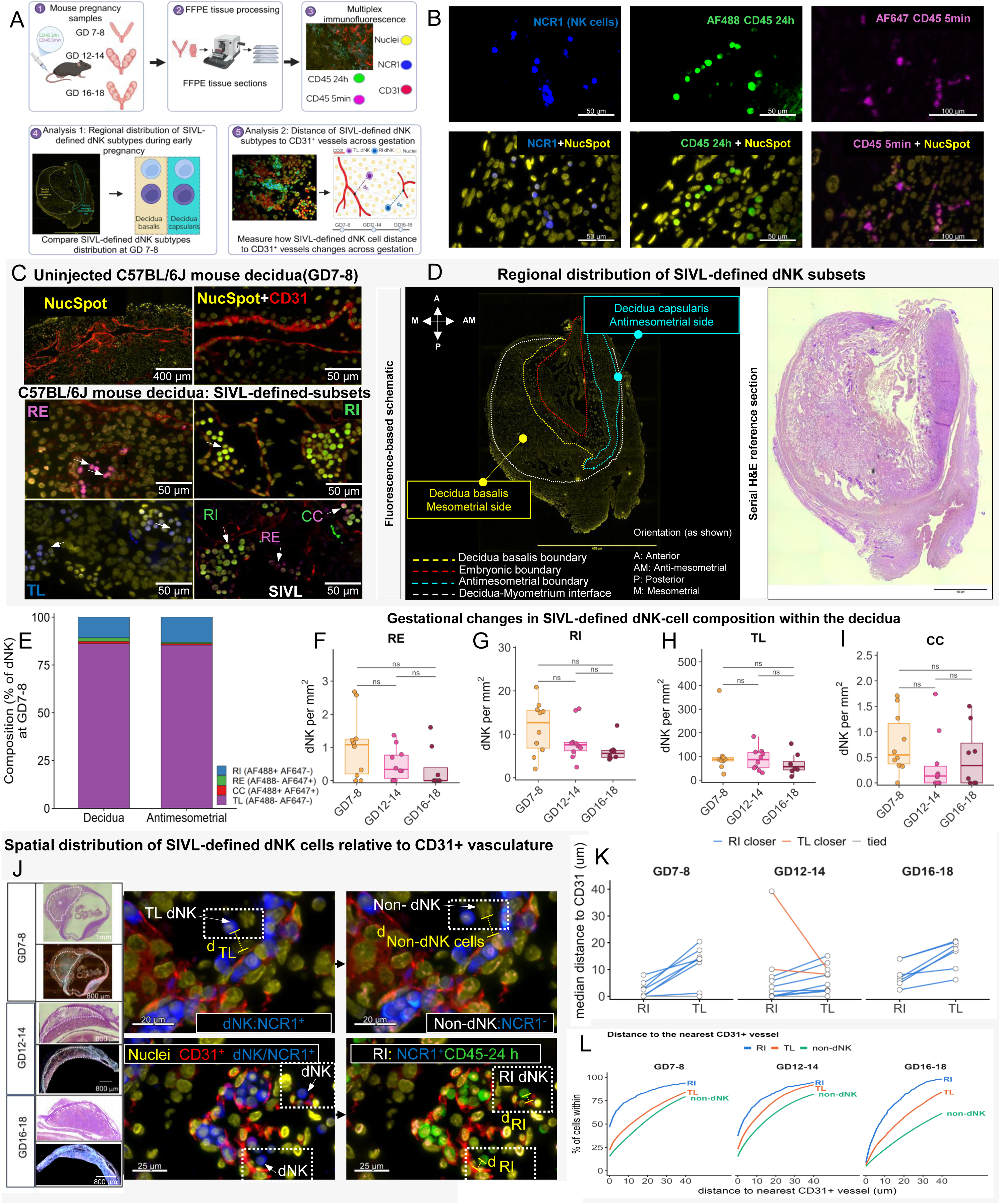
Spatial imaging validate SIVL-defined dNK compartments and reveals compartment-specific perivascular localization across pregnancy. **(A)**Experimental and analytical workflow. Implantation sites were collected from SIVL-injected C57BL/6J mice at GD7-8, GD12-14, and GD16-18, processed as formalin-fixed, paraffin-embedded tissue, and analyzed by multiplex immunofluorescence. NCR1^+^ dNK cells were classified according to their 24-hr and 5-min intravascular CD45 labels. Analyses compared dNK density and SIVL composition between mesometrial and antimesometrial implantation regions at GD7-8 and quantified the distance of individual dNK cells from the nearest CD31⁺ vessel across gestation. **(B)** Representative single-channel and merged immunofluorescence images showing NCR1, CD45-24 h, CD45-5 min, and nuclear stains (NucSpot) within a decidual section. Arrowheads indicate representative NCR1^+^ cells with corresponding intravascular CD45 signals. Scale bars: 50 µm **(C)** Representative images from uninjected and SIVL-injected mouse decidua illustrating background fluorescence, intravascular CD45 labeling, NCR1, CD31⁺ vasculature, and the assignment of SIVL-defined compartments. **(D)** Anatomical delineation of the decidua basalis on the mesometrial side and decidua capsularis on the antimesometrial side in a immunofluorescence image and corresponding serial H&Estained reference section. **(E)** Mean relative composition of TL, RI, RE, and CC dNK subsets within the mesometrial and antimesometrial regions at GD7-8, normalized to 100% of SIVL-classified dNK cells. **(F-I)** Densities of CC, RE, TL and RI dNK cells across gestational ages, shown as cells per mm². Boxplots show medians and interquartile ranges, and individual points represent mice. gestational differences were assessed using the Kruskal-Wallis test, with pairwise comparisons indicated by horizontal bar; ns, not significant **(J)** Representative immunofluorescence images of mouse decidua. Upper: Paired views of the same field showing distances from NCR1⁺ dNK and NCR1⁻ non-dNK cells to the nearest CD31⁺ vessel (red). Lower: Paired views of the same showing the same RI dNK cells positive for both NCR1 (blue) and AF488 (green), with distances measured to the nearest CD31⁺ vessel (red). Yellow brackets indicate shortest cell-to-vessel distances. Scale bars: 20 µm (upper) and 25 µm (lower; images are captured from decidua at GD12-14). **(K)** The distance to the nearest CD31⁺ vessel for each RI and TL NCR1⁺ cells was measured, and the median distance was calculated separately for RI and TL cells within each mouse. Each line connects the paired RI and TL median distances from one mouse at the three gestational stages. Overall paired comparisons across gestational stages were significant by a two-sided sign test (p = 6.6 × 10⁻⁵) and Wilcoxon matched-pairs signed-rank test (p = 0.0007). **(L)** Cumulative distributions of the distance from RI cells, TL cells, and non-dNK cells to the nearest CD31⁺ vessel at each gestational stage. The x-axis indicates per-cell distance to the nearest CD31⁺ vasculature, and the y-axis indicates the cumulative percentage of cells within the specified distance. The number of mice per group is provided in Table 1.

We first compared dNK cell density between the two decidual regions. Paired analysis revealed significantly more dNK cells per mm² in the mesometrial decidua basalis than in the antimesometrial decidua capsularis (median ∼100 vs ∼30 dNK cells per mm²; p=0.023, n=8 mice; supplementary Fig. S2B). Next, we compared the proportional distribution of SIVL-defined dNK compartments between the two decidual regions. In both regions, the TL compartment constituted more than 80% of dNK cells, followed by RI cells (∼15%), whereas RE and CC cells represented only minor fractions (∼1% and ∼2%, respectively) (Fig. 3E and Supplementary Fig. S2C). Thus, the higher dNK-cell density in the decidua basalis was not accompanied by significant regional differences in the proportions of SIVL-defined dNK subsets.

SIVL-defined dNK-cell density quantification showed that TL cells predominated throughout gestation, while RI-cell density was numerically highest during early pregnancy. Notably, SIVL compartments did not reach significance compared across gestational ages, although consistent with trends seen in aggregate flow data (Fig. 3F-I, Fig.1G, Supplementary Fig. S2D).

We next asked where RI, TL, and non-dNK subsets were located relative to CD31⁺ vessels within the decidua. We quantified vessel-wall association as the percentage of each population whose cells overlapped the segmented CD31⁺ vessel wall (distance = 0 μm). RI enrichment was calculated for each mouse by dividing this percentage in RI cells by the corresponding percentage in non-dNK cells: enrichment = (% of RI cells at distance 0) / (% of non-dNK cells at distance 0, where each percentage has its own class as the denominator. RI contact is RI-at-0 μm ÷ all RI cells and the reference is non-dNK-at-0 μm ÷ all non-dNK cells (supplementary Fig. S2E and F). RI cells were significantly enriched at the vessel wall at GD7-8 and GD12-14, with median enrichment of 2.71-fold and 2.07-fold, p= 0.0039 and p = 0.027, respectively, whereas enrichment at GD16-18 was not significant (1.81-fold, Supplementary Fig.S2G).

We then compared the median distances of RI and TL dNK cells to the nearest CD31⁺ vessel wall across gestation (Fig.3J). Paired analysis within each mouse showed significantly shorter distances for RI than for TL cells (Fig. 3K). Consistently, pooled cumulative distance distributions showed a leftward shift of the RI curve relative to the TL and non-dNK curves, indicating that RI dNK cells maintained closer vascular proximity across pregnancy (Fig. 3L).

### 3.5. In situ proliferation complements trafficking in supporting the dNK-cell pool, while active caspase-3 declines across gestation

Next, we asked whether local proliferation and apoptosis contribute to the gestational remodeling of the SIVL-defined dNK-cell compartment (Supplementary Fig.3A and B). To assess proliferation, mice received an intraperitoneal pulse of BrdU 24 hours before tissue collection (Fig. 1E). BrdU incorporation and active caspase-3 positivity were quantified among viable CD45⁺Lin⁻NK1.1⁺ cells (Fig. 4A). At the virgin baseline, BrdU incorporation was low among adult virgin uterine NK cells, averaging approximately 0.4%, compared with 3.1% in the adult spleen and 3.8% in Peyer’s patches; however, the overall baseline comparison did not reach statistical significance (Welch ANOVA, (p=0.06); Fig. 4B and Supplementary Fig.S3A).

**Figure 4.**
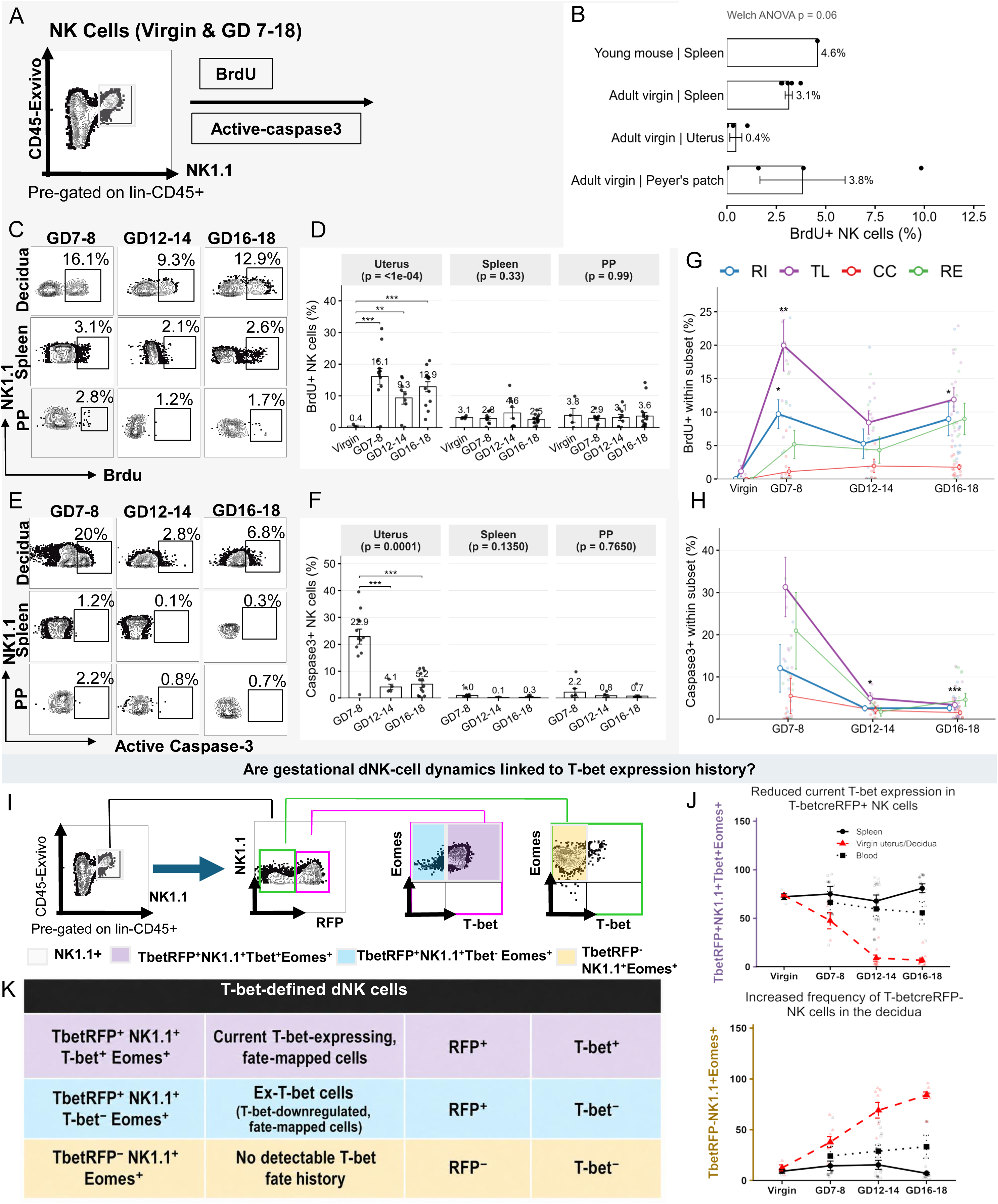
Gestational regulation of BrdU incorporation and active caspase-3 across SIVL- and T-bet-defined uterine/decidual NK-cell populations. (A) Representative gating strategy used to quantify BrdU uptake and active caspase-3 positivity within Lin⁻CD45⁺NK1.1⁺ cells. (B) Baseline frequencies of BrdU⁺ NK cells in the spleen of a young mouse and in the spleen, uterus, and Peyer’s patches (PP) of adult virgin mice. Bars show mean ± SEM; the overall P value was determined by Welch’s ANOVA. (C) Representative flow-cytometric plots showing BrdU incorporation in decidual, splenic, and PP NK cells across pregnancy. Numbers indicate the percentage of BrdU⁺ decidual NK cells. (D) Frequencies of BrdU⁺ NK cells in the uterus/decidua, spleen, and PP at virgin and across gestation. Overall P values are displayed for each tissue. (E) Representative plots showing active caspase-3 positivity in decidual, splenic, and PP NK cells across gestation. (F) Frequencies of active caspase-3⁺ NK cells in the uterus/decidua, spleen, and PP across gestation. Statistical analysis used Welch’s analysis of variance compared virgin tissues (B) gestational stages within each tissue (D, F), with Games–Howell post hoc comparisons in D and F. (G) BrdU incorporation within TL, RI, RE, and CC uterine/decidual NK-cell subsets across gestational stages. (H) Active caspase-3 positivity within the SIVL-defined uterine/decidual NK-cell compartments across gestation Statistical analysis for (G,H) used linear mixed models included gestational stage, trafficking subset, and their interaction as fixed effects, a random intercept per mouse, and subset-specific residual variances. Stage × subset interactions were assessed using joint tests. Tukey-adjusted stage contrasts were performed within each subset; stars indicate comparisons with virgin mice (G) or GD7-8 (H). *P < 0.05, **P < 0.01, ***P < 0.001. (I) Representative gating strategy used to distinguish TbetRFP⁺T-bet⁺Eomes⁺ cells with current T-bet expression, TbetRFP⁺T-bet⁻Eomes⁺ ex-T-bet cells with a history of T-bet expression, and TbetRFP⁻Eomes⁺ non-fate-mapped cells within the Lin⁻CD45⁺NK1.1⁺ population. **(J)** Frequencies of current T-bet⁺ cells within the TbetRFP⁺Eomes⁺ fate-mapped population (upper panel) and TbetRFP⁻Eomes⁺ (lower panel) within the NK-cell population in the uterus/decidua, spleen, and blood at virgin and throughout pregnancy. **(K)** Summary table defining NK1.1⁺Eomes⁺ subpopulations by T-bet fate-mapping status and current expression: TbetRFP⁺T-bet⁺Eomes⁺ (RFP⁺T-bet⁺), TbetRFP⁺T-bet⁻Eomes⁺ (RFP⁺T-bet⁻), and TbetRFP⁻Eomes⁺ (RFP⁻Tbet-).

Pregnancy was associated with a marked increase in BrdU incorporation among dNK (overall p < 0.0001; Fig. 4C and D). BrdU-positive fraction increased from 0.4% in virgin uterine NK cells to approximately 16% at GD7-8 and remained elevated at mid-to-late gestation, whereas no significant gestational variation was detected among splenic NK cells (p=0.33) or Peyer’s patch NK cells (p=0.99; Fig.4D and Supplementary Fig. S4C-D). Active caspase-3 positivity also varied across gestation and peaked at GD7-8 (approximately 23% of dNK cells were active caspase-3 positive) when BrdU incorporation was numerically highest, then declined markedly at subsequent gestational stages. In contrast, active caspase-3-positive NK cells remained infrequent in the spleen and Peyer’s patches (Fig. 4E, F and Supplementary Fig. S4F).

Does trafficking status (SIVL compartment) impact dNK proliferation and apoptosis? BrdU-uptake quantitation stratified by SIVL (RI, TL, CC and RE) revealed that TL population showed the highest within-subset BrdU uptake throughout pregnancy, reaching approximately 20% at GD7-8 before settling closer to 10% (GD12-14, GD 16-18; Fig 4G). Notably, dNK cells which have had vascular exposure (CC, RI, and RE dNK cells) remained comparatively lower. Apoptosis susceptibility was broadly similar to proliferative activity at GD7-8, active caspase-3 positivity was numerically highest among TL and RE dNK cells, with lower frequencies among RI and CC subsets. Unlike proliferation, however, active caspase-3 was far more restrained after the first pregnancy timepoint across all four SIVL-defined compartments (Fig 4E and H). Together, these findings demonstrate that pregnancy-induced dNK-cell remodeling involves gestationally regulated changes in vascular accessibility, proliferative activity, and apoptosis. The compartment-specific distributions of BrdU⁺ and active caspase-3⁺ cells across SIVL-defined dNK cells further indicate that these integrated cellular processes are differentially organized across the dNK-cell pool. We therefore asked whether these integrated cellular were associated with the progressive gestational shift toward T-bet-low dNK populations^36^.

### 3.6. T-bet fate mapping links Eomes^+^ dNK-cell remodeling to differential trafficking and proliferation

We previously showed that T-bet expression is progressively and selectively downregulated in dNK cells across pregnancy^36^, a transcription factor which is involved in egress/trafficking, differentiation, and maturation of both NK cells in a variety of tissues^33,39,40^. Consequently, how current or prior history of T-bet expression is related to gestational remodeling of dNK cells distribution and local proliferation was determined. To address this question, we used B6.RosaRFP/TbetCre fate mapper mice we previously generated ^36^. CD45+Lin^neg^NK1.1⁺ dNK cells were further resolved into three Eomes⁺ populations^35^ based on RFP fate labeling and current T-bet expression: RFP^+^NK1.1^+^Tbet^+^Eomes^+^cells with current T-bet expression, RFP^+^NK1.1^+^Tbet^-^Eomes^+^ ex-T-bet cells with a history of T-bet expression, and RFP^-^NK1.1^+^Eomes^+^ non-fate-mapped dNK cells (Fig. 4I). As previously demonstrated^36^, RFP⁺T-bet⁺Eomes⁺ NK cells progressively and selectively declined in the uterus/decidua across pregnancy, whereas this population remained comparatively stable in the spleen and blood (Fig. 4J, upper panel). Conversely, RFP⁻Eomes⁺ NK cells progressively increased in the decidua from virgin through late gestation, without a comparable increase in the peripheral compartments (Fig. 4J, lower panel).

To determine whether preferential proliferation within the decidua accompanies the remodeling of T-bet-defined dNK states, we compared BrdU uptake among the three states across gestation (Fig. 5A, B and Supplementary Fig. S3G). At GD7-8, approximately 20% of each T-bet-defined dNK population incorporated BrdU. Thereafter, BrdU incorporation was comparatively preserved among dNK cells lacking current T-bet expression, whereas it declined in the current T-bet-expressing dNK subset. BrdU incorporation declined significantly in RFP^+^NK1.1^+^Tbet^+^Eomes^+^ cells and remained low through late gestation. In contrast, both RFP^+^NK1.1^+^Tbet^-^Eomes^+^ andRFP^-^NK1.1^+^Eomes^+^ populations lacking current T-bet expression maintained numerically higher BrdU incorporation during mid-to-late gestation. Similarly, active caspase-3 positivity was highest at GD7-8 across all three T-bet-defined dNK subsets (Fig. 5C and Supplementary Fig. S3H). Positivity subsequently decreased markedly in both fate-mapped populations and remained low through late gestation. In contrast, RFP^-^NK1.1^+^Eomes^+^ cells exhibited a more modest, transient reduction at GD12-14, followed by a partial increase at GD16-18. Thus, the gestational decline in current T-bet-expressing dNK cells was not accompanied by preferential active caspase-3 positivity, suggesting that selective executioner-caspase activation is unlikely to be its primary driver.

**Figure 5.**
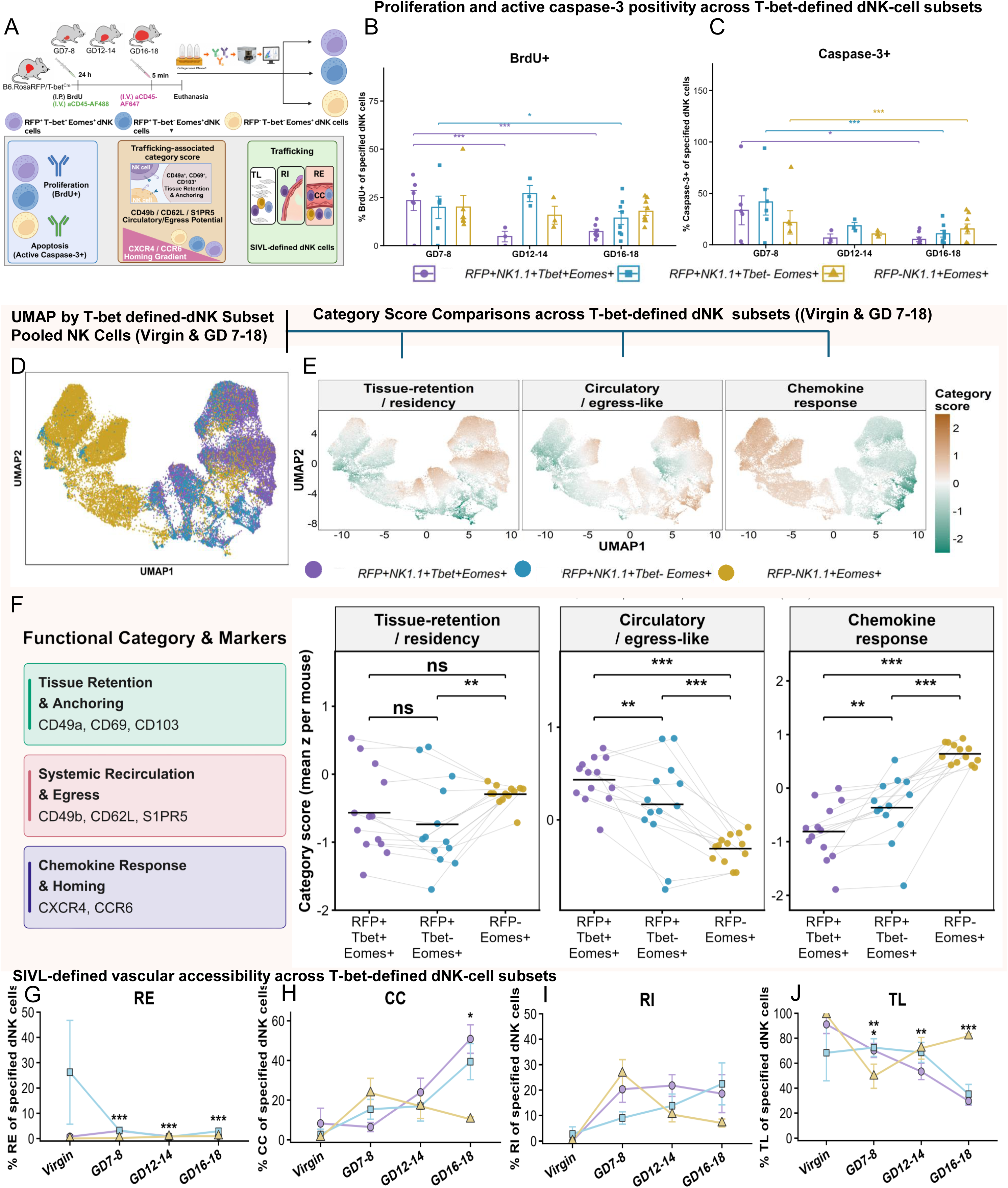
T-bet expression history is associated with dNK-cell proliferation, apoptosis-associated activity, trafficking-related phenotypes, and SIVL-defined vascular accessibility. **(A)** Workflow for integrated analysis of T-bet-defined dNK-cell states. Rosa/Tbet^Cre^ mice were analyzed at three gestational ages following BrdU (I.P.) and anti-CD45-AF488 (I.V.) administration 24 h before euthanasia and anti-CD45-AF647 (I.V.) 5 min before euthanasia. Spectral flow cytometry was used to distinguished Eomes⁺ dNK subsets expressing RFP⁺T-bet⁺, RFP⁺T-bet⁻, or RFP⁻T-bet⁻. Each subset was assessed for BrdU uptake, active caspase-3 positivity, trafficking-associated category scores, and SIVL-defined compartment distribution (TL, RI, RE, and CC).**(B)** Frequencies of BrdU incorporation within the three T-bet-defined Eomes⁺ dNK populations across three gestational stages. **(C)** Active caspase-3 positivity within the corresponding T-bet-defined dNK populations across gestation. individual points represent mice and summary values are presented as mean ± SEM. (B, C): Comparisons within each subset, every gestational age was contrasted against GD7-8 using estimated marginal means with Holm adjustment. Brackets indicate the compared ages; \**P* < 0.05, \*\**P* < 0.01, *** *P* < 0.001. Non-significant comparisons are not added. Statistical analyses were performed using a beta-binomial generalised linear mixed model fitted to positive-cell counts and subset parent totals. **(D)** UMAP distribution of the three T-bet-defined Eomes⁺ dNK populations: **(E)** UMAP distributions of tissue-retention/residency, circulatory/egress-like, and chemokine-response category scores among uterine/decidual NK cells from T-bet fate-mapping mice (Rosa/Tbet^Cre^mice). Brown and cyan indicate higher and lower category scores, respectively. **(F)** Mouse-level mean tissue-retention/residency, circulatory/egress-like, and chemokine-response scores for each T-bet-defined population. Each point represents one mouse (*n* = 13), gray lines connect populations from the same mouse, and horizontal bars indicate group means. Differences among populations were evaluated using linear mixed-effects models followed by estimated-marginal-means comparisons with Holm adjustment. **(G-J)** Proportions of each T-bet-defined dNK population assigned to the (RE; **G**), (CC; **H**), (RI; **I**), and (TL; **J**) subset in the virgin uterus and across gestation. Data are presented as mean ± SEM, with individual points representing mice. Purple circles indicate RFP⁺T-bet⁺Eomes⁺ cells, blue squares denote RFP⁺T-bet⁻Eomes⁺, and yellow triangles denote TbetRFP⁻Eomes⁺ cells. Statistical analysis was performed using Beta-binomial generalized linear mixed model; ** *P* < 0.01 and *** *P* < 0.001; ns, not significant. The number of mice per group is provided in Table 1.

Next, we asked whether the three T-bet-defined dNK states differed in their tissue-retention/residency, circulatory/egress-like, and chemokine-response potential. To address this, we overlaid the RFP/T-bet-defined states onto the shared phenotypic UMAP using the same eight trafficking-associated markers used to calculate the category scores: CD49a, CD69, and CD103 for tissue retention/residency; CD49b, CD62L, and S1PR5 for circulatory/egress-like potential; and CXCR4 and CCR6 for chemokine responsiveness (Fig.5D). UMAP visualization showed partial phenotypic separation among the three T-bet-defined NK populations along continuous trafficking-associated gradients rather than into discrete clusters. RFP⁻NK1.1⁺Eomes⁺ subset displayed a chemokine-responsive, low-egress profile, characterized by higher chemokine-response and lower circulatory/egress-like scores. In comparison, RFP⁺NK1.1⁺T-bet⁺Eomes⁺ cells and RFP⁺NK1.1⁺T-bet⁻Eomes⁺ ex-T-bet cells displayed substantial phenotypic overlap, although cells retaining T-bet expression were relatively shifted toward higher circulatory/egress-like scores (Fig. 5E). For quantitative comparison, across pooled gestational ages, cell-level category scores were averaged within each T-bet-defined state for each mouse, then analyzed using a linear mixed-effects model with T-bet-defined state as a fixed effect and mouse as a random intercept, followed by Holm-adjusted pairwise comparisons of estimated marginal means (Fig. 5F). The tissue-retention/residency score was significantly higher in RFP^-^NK1.1^+^Eomes^+^ non-fate-mapped cells than in RFP^+^NK1.1^+^Tbet^-^Eomes^+^ dNK cells, whereas comparisons involving RFP^+^NK1.1^+^Tbet^+^Eomes^+^ cells were not significant. Interestingly, the circulatory/egress-like score was highest in RFP^+^NK1.1^+^Tbet^+^Eomes^+^ cells, intermediate in ex-T-bet cells, and lowest in RFP^-^NK1.1^+^Eomes^+^ dNK cells. Conversely, the chemokine-response score was lowest in RFP^+^NK1.1^+^Tbet^+^Eomes^+^ cells, intermediate in ex-T-bet cells, and highest in RFP^-^NK1.1^+^Eomes^+^ dNK cells.

We next asked if vascular accessibility is modified in Tbet-defined dNKs across pregnancy (Fig. 5G-J). Among both RFP^+^NK1.1^+^Tbet^+^Eomes^+^ and RFP^+^NK1.1^+^Tbet^-^Eomes^+^ dNK cells, the tissue-localized (TL) fraction declined significantly between the virgin uterus and GD16-18, whereas the continuously circulating (CC) fraction increased significantly over the same interval. In contrast, RFP^-^NK1.1^+^Eomes^+^ non-fate-mapped dNK cells exhibited a significant but transient reduction in tissue localization at GD7-8, followed by recovery at mid-and late gestation; by GD16-18, these cells remained predominantly tissue localized (TL), with only a minor continuously circulating fraction (CC). The RI compartment remained comparatively modest and did not exhibit a consistent gestational pattern across the three RFP/T-bet-defined dNK states.

Collectively, these findings reveal distinct gestational stage-specific trajectories among T-bet-defined dNK states: fate-mapped populations became increasingly represented in the continuously circulating compartment during late gestation, whereas T-bet^cre^RFP^-^ NK1.1^+^Eomes^+^ cells remained predominantly tissue-localized, and proliferative activity was comparatively preserved among cells lacking current T-bet expression. Thus, these findings support a model in which the integrated processes of selective trafficking and local proliferation contribute to a shift in the dNK-cell pool from a T-bet-high to a T-bet-low phenotype.

## 4. Discussion

A central finding of this study is that phenotypic remodeling of the dNK-cell compartment begins early in pregnancy, marked by the decidual enrichment of CD49a⁺CD49b⁺ dNK cells. The co-expression of CD49a and CD49b identifies an early pregnancy dNK state that falls outside the conventional binary classification of CD49a⁺CD49b⁻ trNK and CD49a⁻CD49b⁺ cNK cells^41^. The early accumulation of this population may represent a gestationally induced dNK-cell state undergoing phenotypic adaptation within the decidual environment. Consistently, a previous study identified a uterus-restricted Eomes⁺CD49a⁺CD49b⁺ NK subset enriched for the immature CD11b^low^CD27^high^ phenotype and capable of producing both IFN-γ and TNF^19^. Our finding supports a model in which recruited CD49b⁺ cNK cells progressively upregulate CD49a and downregulate CD49b, while retaining Eomes expression after entering the decidua (bioRxiv)^42^. Nevertheless, the selective enrichment of CD49a⁺CD49b⁺ dNK cells during early pregnancy highlights the limitations of inferring tissue residency from CD49a and CD49b expression alone and supports a model in which dNK-cell phenotype and localization are dynamically remodeled across pregnancy.

We employed SIVL to carefully delineate the features of dNK cells entering the uterus across gestation. In the virgin uterus, the predominance of the TL dNK population was consistent with previous evidence that the nonpregnant murine uterus is enriched in CD49a⁺-defined trNK cells, whereas CD49a⁻-defined cNK cells are comparatively low^5^. However, SIVL analysis revealed that pregnancy does not merely expand the pre-existing TL dNK-cell pool. At GD7-8, the presence of RI dNK cells indicated NK-cell entry from the circulation, while the concurrent CC compartment identified a distinct population that remained vascularly accessible (either fully in the vasculature or adherent to the surfaces of intervillous space). Thus, early recruitment of circulating NK cells contributes to dNK-cell accumulation alongside the persistence of a substantial tissue-localized population, extending our previous findings indicating that decidual leukocyte trafficking remains active throughout pregnancy^7^. As pregnancy advanced, the increasing representation of the CC dNK compartment was concordant with the expansion of the CD49b⁺ cNK cells, suggesting a progressively greater vascular contribution; however, CC status should not be considered equivalent to a cNK phenotype^20,43^. Rather, the dual-labeled CC compartment may include freely circulating cells captured at euthanasia as well as cells loosely or firmly associated with the luminal surfaces of decidual vessels during the 24-hour interval.

Moreover, SIVL revealed that gestational changes in immune cell trafficking were lineage-specific. Compared with CD4⁺ and CD8⁺ T cells, NK cells maintained a pronounced TL fraction throughout gestation, consistent with their sustained association with the decidual stroma^5^. CD4⁺ and CD8⁺ T cells exhibited a greater contribution from the vascularly accessible CC and RI compartments, indicating more extensive exchange with the circulation. These findings provide mechanistic insight into the well-described transition from an NK-cell-dominated decidual immune environment during early pregnancy to greater T-cell representation later in pregnancy. Early in murine pregnancy, decidual stromal cells restrict the recruitment of activated effector T cells through epigenetic silencing of T-cell-attracting chemokines including *Cxcl9*, *Cxcl10*, and *Ccl5* ^44^. However, this restriction is not equivalent to complete exclusion, as our previous SIVL analysis showed increasing proportions of vascularly accessible decidual leukocytes during mid-to-late murine gestation and identified a tissue-localized T-cell population late in pregnancy^7^.

Interestingly, SIVL further resolved dNK cells phenotypic heterogeneity. Its trafficking patterns were accompanied by a distinct phenotypic identity within the dNK-cell compartment. Marker-based category scoring resolved three coordinated yet partially overlapping programs associated with tissue retention, circulatory/egress potential, and chemokine responsiveness. Furthermore, the marker trajectories do not support a fixed binary separation between resident and circulating dNK cells or a linear progression through the four SIVL compartments. Instead, dNK subsets with different vascular-accessibility histories retain mixed phenotypic features, and the molecular basis of tissue localization changes across gestation. The early increase in the tissue-retention-associated score was primarily accompanied by elevated CD49a and CD69, particularly within the TL at GD7-8. This phenotype is consistent with the enrichment of CD49a⁺CD69⁺ dNK populations shown by parabiosis to be locally maintained and with the predominance of CD49a⁺Eomes⁺ trNK cells during early murine pregnancy ^5,20^.

CD49a supports adhesion to collagen-containing extracellular matrices, whereas CD69 is widely associated with tissue localization and can restrict S1PR1-dependent leucocyte egress ^45^. However, this mechanism does not extend to S1PR5-mediated NK-cell trafficking because S1PR5 does not interact with, and is not inhibited by, CD69^33^. Notably, the subsequent reduction in CD49a and CD69 expression was not accompanied by a uniform decline in all retention-associated markers. Instead, CD103 became relatively more prominent during late gestation, indicating a shift in the molecular basis of tissue interaction rather than a simple progressive loss of residency-associated properties^46^. Similarly, circulatory/egress-like features were composed of a nonuniform distribution of CD49b, CD62L, and S1PR5 at different gestational ages, again indicating stage-specific specialization. Conventionally, CD49b is principally associated with conventional NK-cell phenotype, CD62L supports entry into secondary lymphoid tissues, and S1PR5 provides a functional signal for NK-cell egress from the bone marrow and lymph nodes^33,47^. Their nonparallel expression across SIVL-defined compartments indicates that the circulatory/egress score integrates distinct but complementary features of NK-cell trafficking rather than representing a single egress pathway. TL dNK cells showed a gestational reduction in the circulatory/egress-like score, driven largely by decreased CD49b. Although their relatively preserved S1PR5 expression is consistent with retained potential to respond to sphingosine-1-phosphate gradients, receptor expression alone does not mandate functional signaling or active egress from the decidua. In contrast, RI dNK cells displayed the highest circulatory/egress-like score among the pregnant compartments at mid-to-late gestation. Notably, their relative maintenance of CD49b and CD62L, despite lower S1PR5 expression, is consistent with a transitional state in which recently recruited NK cells retain circulation-associated features after decidual entry while progressively acquiring a tissue-resident phenotype(bioRxiv) ^42^. CC dNK cells retained relatively high CD49b but exhibited lower CD62L and S1PR5, supporting a predominantly conventional NK-like state rather than uniformly elevated egress competence.

In contrast to the early peak in the retention-associated score, the chemokine-response score generally increased from the virgin uterus toward mid-to-late pregnancy. The heatmap indicated that this trajectory was associated with increased CXCR4 and CCR6 expression, most prominently within RI and RE dNK compartments. CXCR4 is directly relevant to NK-cell positioning at the maternal-fetal interface because decidual and trophoblast-derived CXCL12 can attract CXCR4-expressing dNK cells^16^. In a mouse study, adult-induced global maternal Cxcr4 deletion reduced dNK-cell representation, disrupted their spatial distribution and granzyme-B expression, while transplantation of CXCR4-sufficient bone marrow substantially rescued these abnormalities^11^. Complementary evidence showed that adoptive transfer of CXCR4⁺ NK cells into pregnant Nfil3⁻/⁻ mice reduced embryo resorption and restored the decidual Th2/Th1 balance^48^. Recently, a human study characterized CXCR4^+^CD56bright dNK cells as a less cytotoxic, IL-4-producing immunoregulatory population that was reduced and functionally impaired in recurrent miscarriage^48^. Moreover, reduced CXCR4⁺ dNK-cell abundance in recurrent spontaneous abortion has also been linked to impaired trophoblast autophagy through an IGF2-TP53-CXCR4 regulatory pathway^49^. CCR6 is another component of the gestationally enhanced chemokine-response program; however, unlike the CXCL12-CXCR4 axis, its contribution to dNK-cell recruitment and positioning at the maternal-fetal interface remains poorly defined. CCR6 recognizes CCL20 and may identify responsiveness to inflammatory or mucosal chemotactic signals. The gestational increase in CCR6 across the SIVL-defined compartments, particularly RE and CC populations during mid-to-late gestation, posits a possibility of an increased capacity to respond to CCL20, a hypothesis we have not directly tested. In the virgin uterus, CCR6 is predominantly expressed on CD127⁺RORγt⁺ LTi-like/ILC3 populations rather than Eomes⁺ NK cells^50^. In murine pregnancy, evidence for the CCL20-CCR6 axis derives primarily from other immune populations: CCR6 expression on circulating and decidual Tregs increased during allogeneic compared with syngeneic pregnancy, together with direct migration of purified Tregs toward CCL20 in vitro^51^. In humans, first-trimester CD56⁺CD16⁻ dNK cells expressed CCR6, although their migration toward CCL20 was not examined^16^. Moreover, decidual NCR⁺ ILC3 and LTi-like cells expressed higher levels of CCR6 than mature NK cells ^52^. Clinical studies further associate dysregulation of the CCL20-CCR6 axis, with reduced peripheral blood and decidual CCL20 abundance and CCR6^+^ regulatory T cells in recurrent miscarriage, but increased circulating CCL20 in preeclampsia^53,54^. These findings imply enhanced potential to respond to locally produced chemokines as pregnancy progresses, but it does not distinguish active migration from gestational induction of receptor expression or selective enrichment of receptor-positive cells. Importantly, the relationship between these programs and the SIVL compartments showed that vascular accessibility and trafficking phenotype were related but not interchangeable. CC dNK compartment exhibited a lower tissue-retention-associated score than the TL, RI, and RE compartments, supporting an association between sustained vascular accessibility and reduced expression of tissue-localization markers. However, the remaining programs did not follow a simple linear transition from circulating to resident states. RI cells exhibited greater circulatory/egress-like features than RE and CC cells, while RE cells displayed greater chemokine responsiveness than RI subset.

Spatial mapping of SIVL-defined dNK populations further clarified the anatomical pattern of their vascular-accessibility. The greater dNK-cell density in the decidua basalis was consistent with the mesometrial accumulation of uNK cells previously described during early mouse gestation^55,56^. However, the absence of preferential enrichment of any particular SIVL compartment between the decidua basils and decidua capsularis suggests broadly similar dNK vascular-accessibility profile. Furthermore, the predominance of TL across gestation complements previous evidence that uterine trNK cells substantially contribute to dNK-cell accumulation through local proliferation^5,20^, although given design, TL status in this study does not test long-term tissue residency.

A key finding of our spatial analysis was that vascular proximity differed among SIVL-defined dNK subsets. Earlier histological studies documented perivascular and intravascular accumulation of dNK cells in the mouse decidua^14^. Recent spatial transcriptomic studies have identified distinct immune population and vascular microenvironment in the mouse decidua^57^, while human imaging studies have shown NK-cell accumulation around remodeling spiral arteries^58,59^. Our analysis extends this spatial perspective by linking vascular proximity to SIVL-defined vascular-accessibility history, where RI cells were consistently closer to vessels than TL cells across gestation. This distinct vascular association raises the possibility that RI and TL populations differ in their interactions with the vascular microenvironment. Thus, whether the preferential vascular positioning of RI cells reflects recent tissue entry, retention near vessels, or functional specialization remains to be examined.

Importantly, our data clearly demonstrate that abundance of TL dNK compartment is due to a dynamically maintained dNK-cell pool and should not be interpreted as evidence of a static resident population. The gestational peak in decidual BrdU uptake, without a corresponding change in the spleen or Peyer’s patches, supports tissue-specific regulation of local proliferation. This finding is consistent with previous studies using BrdU incorporation and Ki-67 staining that showed intensive NK-cell proliferation within the early mouse decidua without a comparable increase in the spleen^2,5,60^. Nevertheless, while BrdU uptake predominated in TL cells, it was also detected in CC and RI cells, indicating that recent proliferative activity was not wholly restricted to the TL compartment. Similarly, the early gestational peak in active caspase-3 supports this model in which the dNK-cell pool is dynamically maintained. Importantly, apoptotic activity was not restricted to the actively cycling TL compartment but was also detected in RE subset. Importantly, these are not systemic changes as we show that splenic active caspase-3 is unchanged across gestation consistent with prior studies using TUNEL assessment of splenic NK1.1⁺ cells in pregnancy^61^. Earlier studies revealed uNK-cell DNA fragmentation and loss during late gestation^62^, whereas we observed an early gestational peak in active caspase-3, however, that was in isolated, cultured anti-Fas treated uNK cells, not in situ^62^. Transient caspase-3 activation during proliferation may reflect regulation of activated or proliferating dNK-cell states that precedes the definitive loss of mature cells^63^ later in pregnancy, however this mechanism remains to be examined. Consistently, recent study demonstrated that declining proliferative activity was accompanied by the emergence of a terminal differentiation-associated apoptotic program^60^. In humans, increased apoptotic activity within the decidua was associated with recurrent miscarriage however it predominated in decidual stroma rather than immune cells^64,65^. Rhesus macaque studies have examined apoptosis at the MFI, but have not resolved apoptotic activity specifically within dNK cells^66,67^. Collectively, our data supports a model in which integrated processes of local proliferation, apoptosis and trafficking dynamically maintain dNK-cell residency across gestation rather than representing it as a fixed cellular state.

Understanding the transcriptional regulation of this dynamic residency process requires distinguishing their roles in NK-cell development from their contribution in gestational adaptation. dNK cells are regulated by the T-box transcription factors T-bet and Eomes, each act at distinct checkpoint in dNK-cell development and differentiation^29,31^. Conditional deletion of Eomes in Ncr1-expressing cells eliminated uterine CD49a^+^NK cells in virgin and pregnant mice^35^, whereas CD49a⁺ uterine NK cells remained detectable in globally T-bet-deficient virgin mice^68^. Our previous finding reinforces this distinction: dNK cells progressively and selectively downregulate T-bet across gestation while retaining Eomes expression^36^. This transition was not accompanied by loss of NK-cell identity but was associated with increased expression of homing-and tissue-residency-associated genes, reduced cytotoxic programs, and mid-gestational suppression of IFN-γ signaling^36^. These findings support the hypothesis that T-bet modulation contributes to this gestational dynamic within an Eomes-expressing dNK-cell population.

Proliferative activity provides a potential link between this gestational T-bet downregulation and the maintenance of the dNK-cell pool. Although the early gestational proliferative peak did not preferentially involve a particular fate-mapping subset, proliferation was subsequently sustained preferentially in ex-T-bet and non-fate-mapped RFP⁻ dNK populations. These findings complement previous evidence that the numbers and morphology of dNK populations are maintained despite absent T-bet expression^69^. In addition, this redistribution of proliferative activity toward T-bet-low states may parallel the uNK-cell differentiation program described by Han et al., which is initiated by IL-21R– STAT3 signaling and accompanied by progressive *Tbx21* downregulation. Consistently, studies in mouse spleen and bone marrow demonstrated increased proliferation in mature NK-cell subsets following *Tbx21* deletion^29,60^. In contrast, apoptosis did not correlate with T-bet expression, as neither its early gestational peak nor its subsequent decline preferentially involved a particular T-bet-defined subset.

The category-score analysis further suggests that the gestational shift toward T-bet-low dNK states may involve differences in trafficking and homing potential alongside proliferative maintenance. Although each category marker has distinct and context-dependent functions, their integrated expression profiles provide a framework for comparing homing, residency and egress-associated phenotypes. The highest chemokine-responsiveness scores in non-fate mapped RFP⁻ Eomes^+^ dNK cells were consistent with a phenotype favoring chemokine-guided recruitment and positioning within the decidua^48,70^. Conversely, RFP⁺T-bet⁺ Eomes^+^ dNK cells exhibited the highest circulatory/egress scores, suggesting a greater propensity for ongoing trafficking between the circulation and decidual tissue, which may contribute to their changing representation across gestation. This circulatory/egress feature associated with RFP^+^T-bet^+^ Eomes^+^ dNK cells may be mechanistically based in the T-bet-dependent regulation of NK-cell trafficking. T-bet promotes S1PR5 expression and S1PR5-dependent egress from bone marrow and lymph nodes, while T-bet deficiency impairs NK-cell movement from bone-marrow parenchyma into vascular sinusoids^33,71^. However, the markers contributing to this category scores are not uniformly regulated by T-bet. Eomes enhances the acquisition and maintenance of CD49b expression, whereas T-bet deficiency increases CD62L expression in developing bone-marrow NK cells^31,71^. Within the decidua, shared microenvironmental signals may maintain residency-associated phenotypes across T-bet-defined states, potentially explaining the absence of significant differences in tissue-retention scores among the three subsets. Uterine CD49a⁺ NK cells are preserved in T-bet-deficient mice, while TGF-β signaling promotes the conversion of circulating CD49b^+^ NK cells into CD49a^+^ NK cells upon entry into the decidua (bioRxiv)^42^.

The trafficking-associated differences in T-bet fate mapped dNK states captured by the category scores were reflected in the vascular-accessibility profiles resolved by SIVL. The TL compartment remained predominant among RFP⁻ Eomes^+^ dNK population, although its transient reduction in early gestation coincided with increased representation of CC and RI subsets. Conversely, RFP⁺ Eomes^+^ ex-Tbet cells exhibited a progressive decline in the TL fraction and an increase in the CC fraction, indicating a gestational shift toward greater vascular accessibility. Alongside preferentially maintained proliferation in T-bet-low subsets, this trafficking-associated redistribution may contribute to the remodeling of the dNK-cell pool toward T-bet-low states.

Together, our findings support a model in which dNK-cell residency is not a static property but a dynamically maintained state through stage-dependent changes in trafficking, vascular accessibility, proliferation, and residency-associated phenotypes. These integrated processes shape dNK-cell heterogeneity and contribute to the gestational remodeling of T-bet-defined states. The mechanisms linking T-bet adaptation to these processes and their disturbance in disorders of placentation and pregnancy will be the object of further investigation.

## ACKNOWLEDGMENTS

We would like to thank D. Sheerar from UWCCC Flow lab for technical support.

## Funding

Research support was provided by March of Dimes (5-FY18-541), Burroughs Welcome Fund (1019835), R21AI175753 and R01HL163623 (to AKS); MM was supported by ECEB and R21AI175753 (to AKS); JV was supported by K01AI182448 and L70HD119811. UW Carbone Cancer Center Flow Cytometry core was supported by P30 CA014520, 1S10OD025225-01. MVM acknowledges University of Wisconsin Centennial Scholars Program funding and Building Interdisciplinary Research Careers in Women’s Health (BIRCWH) Training Award (NIAMS 2K12AR084227).

## CONFLICT OF INTEREST

The authors declare no competing interests.

## Data availability

The data supporting the findings in this study are available from the corresponding author upon reasonable request.

## AUTHOR CONTRIBUTIONS

**Mona A Mohamed:** Conceptualization, Methodology, Validation, Formal analysis, Investigation, Data Curation, Writing - Original Draft, Writing - Review & Editing, Visualization.

**Jessica Vazquez**: Conceptualization, Methodology, Validation, Data Curation, Formal analysis.

**Marco Cordero:** Investigation, Data Curation, Formal analysis.

**FL Jaimes:** Investigation.

**Sophia Beaulieu**: Investigation

**Eavan Finnerty**: Investigation

**Xiang Yi Cen**: Investigation

**Roghyah Hasan:** Investigation

**María Virumbrales-Muñoz:** Methodology, Validation, Writing-Review & Editing, Supervision

**Aleksandar K. Stanic**: Conceptualization, Writing - Original Draft, Writing - Review & Editing, Supervision, Project administration, Funding acquisition.

## Notes

### Competing Interest Statement

The authors have declared no competing interest.

