## SupplementalFigures for "Serial intravascular labeling resolves dynamic decidual NK residency and gestational remodeling toward T-bet-Low states"

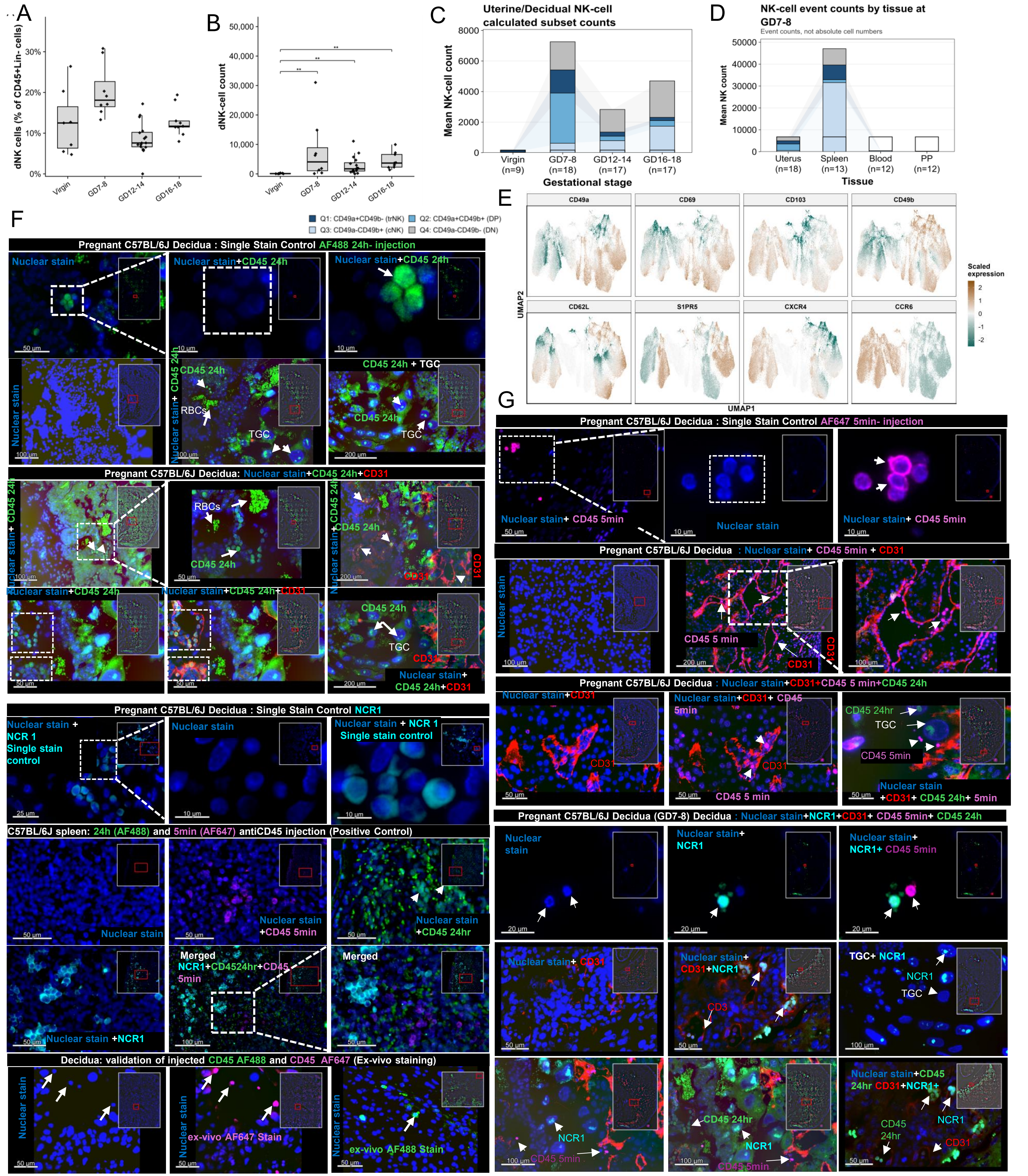


**Supplementary Figure S1. Extended analysis of gestational dNK-cell abundance and validation of fluorescence-based imaging. (A, B)** Frequency of dNK cells among CD45⁺Lin⁻ cells **(A)** and dNK-cell event counts **(B)** in the virgin uterus and decidua at GD7–8, GD12–14, and GD16–18 (virgin, $n$ = 9; GD7–8, $n$ = 18; GD12–14, $n$ = 17; GD16–18, $n$ = 17 mice). Points represent individual mice, and boxes show the median and interquartile range. Statistical comparisons are indicated in the figure. **(C)** Calculated mean counts of CD49a⁺CD49b⁻, CD49a⁺CD49b⁺, CD49a⁻CD49b⁺, and CD49a⁻CD49b⁻ NK-cell populations in the virgin uterus and decidua across gestation. Total bar height represents the mean calculated NK-cell count. **(D)** Mean NK-cell event counts in the uterus/decidua, spleen, blood, and Peyer's patches at GD7–8 (uterus/decidua, $n$ = 18; spleen, $n$ = 13; blood, $n$ = 12; Peyer's patches, $n$ = 12 mice). Values represent acquired events and not absolute tissue cell numbers. **(E)** UMAP feature plots showing scaled expression of CD49a, CD69, CD103, CD49b, CD62L, S1PR5, CXCR4, and CCR6 among uterine/decidual NK cells. Cells from all gestational groups were embedded in a common UMAP (virgin, $n$ = 5; GD7–8, $n$ = 10; GD12–14, $n$ = 16; GD16–18, $n$ = 9 mice; 40 mice total). **(F)** Representative single-channel and merged fluorescence images validating detection of 24-h intravascular anti-CD45–AF488, NCR1–AF350, CD31, and ex vivo CD45 staining. Images include AF488 and NCR1 single-stain controls, a multichannel positive control, and ex vivo staining controls. **(G)** Representative AF647 single-stain and merged multichannel images validating detection and separation of 5-min anti-CD45–AF647, 24-h anti-CD45-AF488, NCR1-AF350, and CD31 labeling. Nuclear staining is shown in blue, NCR1 in cyan, CD31 in red, 24-h anti-CD45–AF488 in green, and 5-min anti-CD45–AF647 in magenta. White arrows indicate representative labeled cells or structures; scale bars are shown within the images. PP, Peyer's patches.

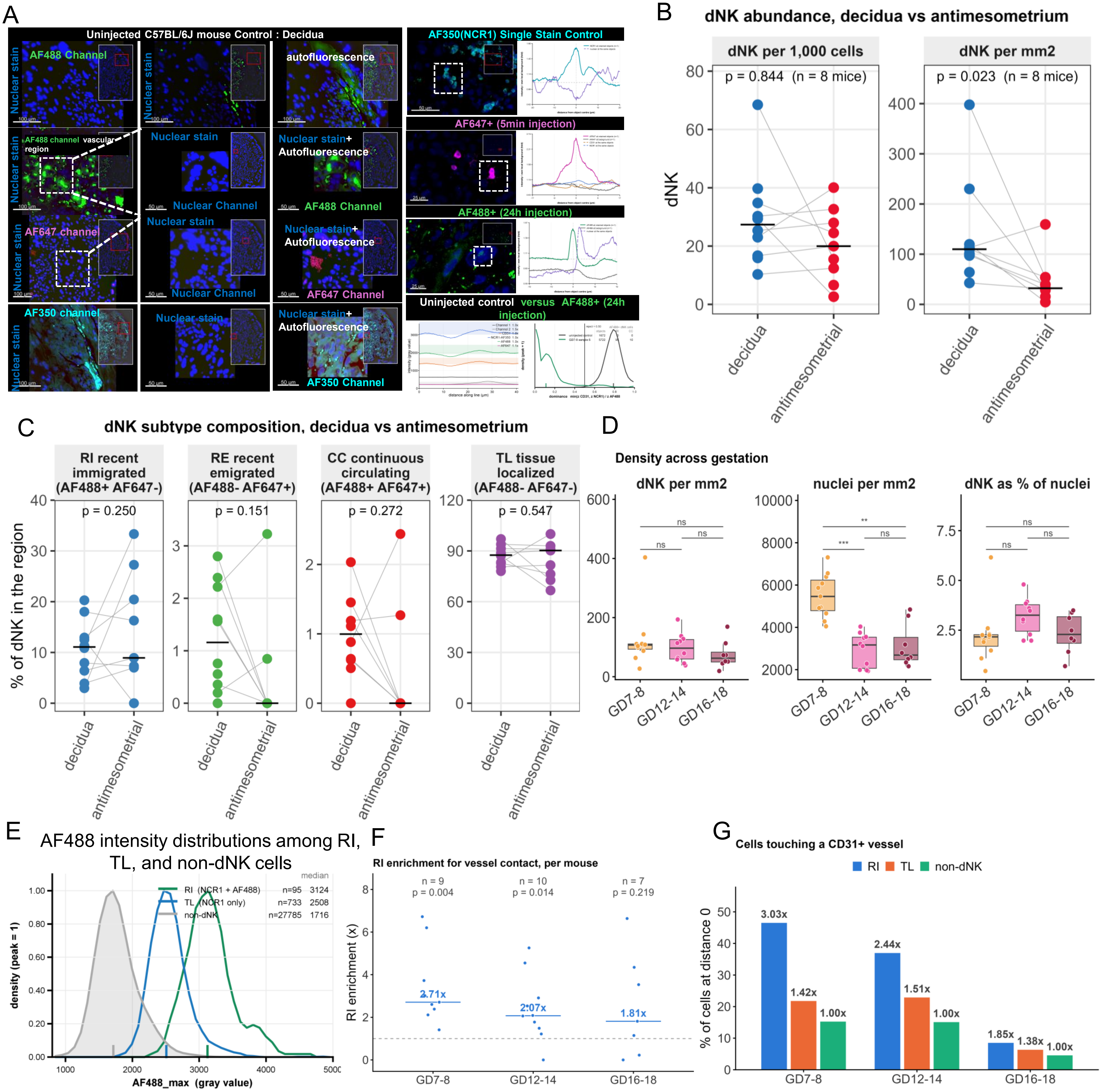


**Figure S2. . Regional distribution and CD31⁺ vessel association of SIVL-defined dNK cells. (A)** Representative single-channel fluorescence images and intensity profiles from an uninjected decidual control, an NCR1–AF350 single-stain control, and tissues collected following 5-min anti-CD45–AF647 or 24-h anti-CD45–AF488 administration.**(B)** Paired comparison of dNK-cell abundance between decidual and antimesometrial regions at GD7-8, expressed as dNK cells per 1,000 total cells and per mm² of tissue. Each point represents one region from an individual mouse; gray lines connect paired regions from the same mouse, and horizontal bars indicate medians. Statistical analysis: paired Wilcxon signed rank test, p=0.844; n=8 mice. **(C)** Paired comparison of the proportions of RI, RE, CC, TL dNK subsets in the decidual and antimesometrial regions at GD7-8. Each point represents the indicated SIVL-defined population as a percentage of total dNK cells from one mouse; lines connect regions from the same mouse, and horizontal bars indicate medians. Statistical analysis: Wilcoxon signed-rank across mice with both regions. **(D)** NCR1⁺ cells, and total nuclei were quantified within annotated decidual regions of FFPE sections at the three gestational stages. dNK density was calculated as the number of NCR1⁺ cells per decidual area (mm²), nuclear density as the total number of nuclei per decidual area, and dNK percentage as NCR1⁺ cells relative to total nuclei. Each point represents one mouse; boxes show the median and interquartile range. Overall gestational differences were assessed using the Kruskal-Wallis test, with pairwise comparisons indicated by horizontal bars. Exact *p* values and significance annotations are shown. ns, not significant; *$p$ < 0.01; ***$p$ < 0.001. **(E)** Distributions of the maximum AF488 intensity among RI dNK cells (NCR1⁺AF488⁺), TL dNK cells (NCR1⁺AF488⁻), and non-dNK cells. Vertical lines indicate group medians; the numbers of analyzed cells and median intensities are reported in the panel. **(F)** Mouse-level enrichment of RI dNK-cell contact with CD31⁺ vessels relative to non-dNK cells from the same mouse. Points represent mice, horizontal bars indicate medians, and the dashed line denotes the null ratio of 1.0. Enrichment was evaluated using a one-sample Wilcoxon signed-rank test against 1.0. **(G)** Percentage of RI, TL, and non-dNK cells in direct contact with CD31⁺ vessels (nearest-vessel distance = 0 μm) across gestation. Values above the bars indicate fold differences relative to the within-stage non-dNK reference, normalized to 1.00. Non-dNK cells were detected nuclei that did not meet the dNK-cell classification criteria.

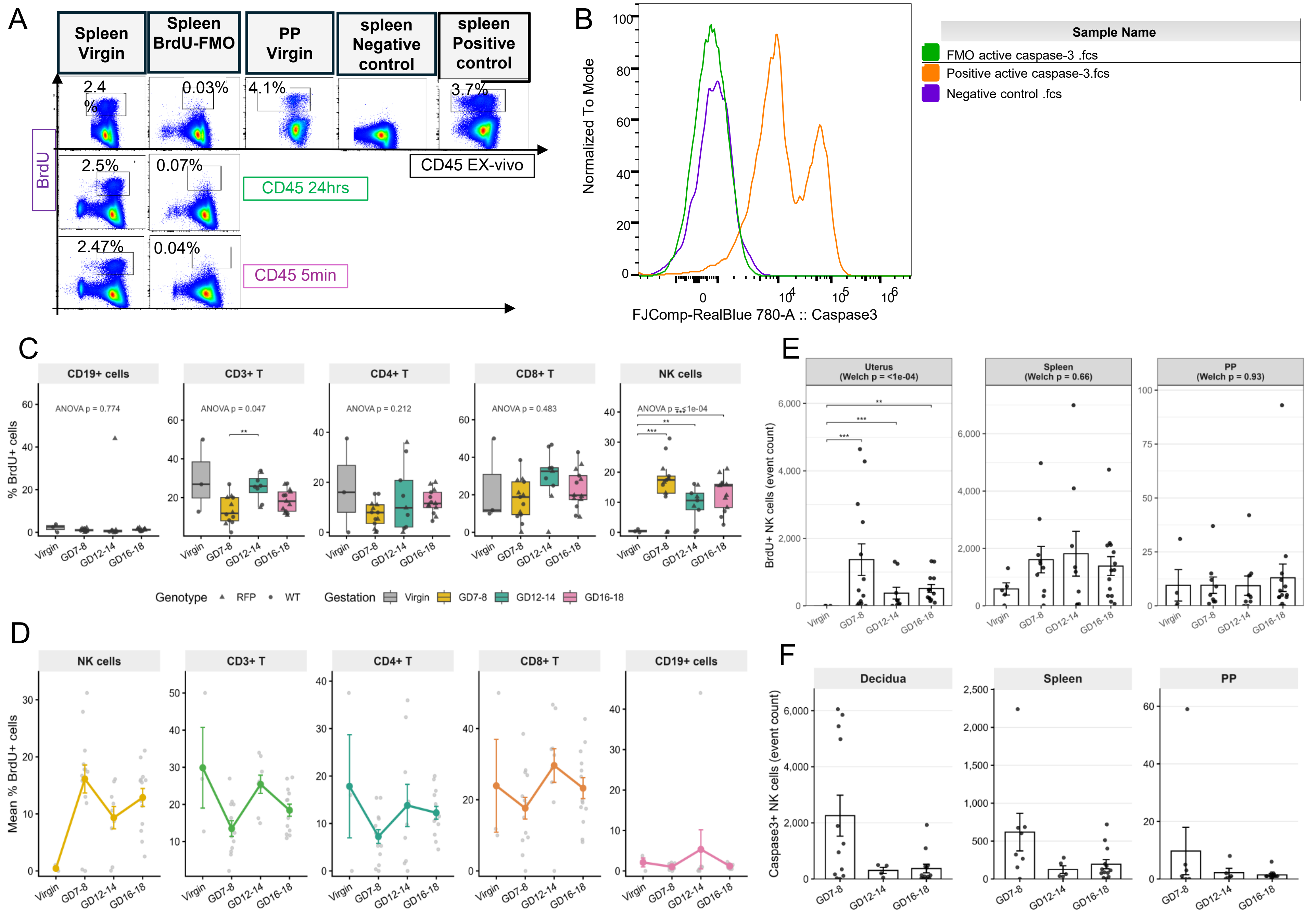


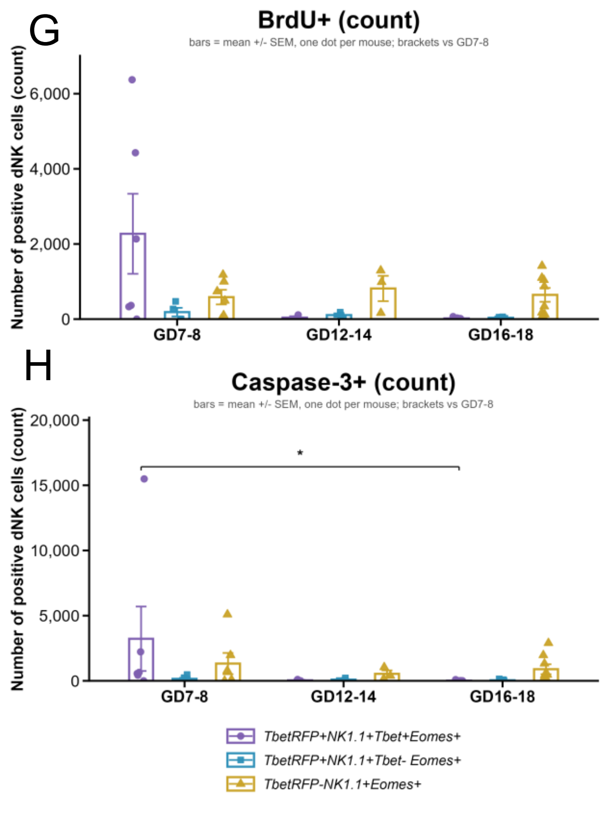


**Supplementary Figure S3. Validation and extended analysis of BrdU incorporation and active caspase-3 across lymphocyte populations and tissues.** **(A)** Representative flow-cytometric plots showing BrdU staining in virgin spleen and Peyer's patches (PP), together with BrdU fluorescence-minus-one (FMO), negative, and positive controls. BrdU staining is shown in relation to ex vivo CD45, 24-h intravascular CD45, and 5-min intravascular CD45 labeling. Numbers indicate the percentage of BrdU$^+$ cells within the corresponding parent population. **(B)** Representative active caspase-3 fluorescence distributions for FMO, negative, and positive controls. **(C)** Percentages of BrdU$^+$ CD19$^+$ cells, total CD3$^+$ T cells, CD4$^+$ T cells, CD8$^+$ T cells, and NK cells in the virgin uterus and decidua across gestation. Points represent individual mice; point shapes distinguish WT and TbetRFP fate-mapper mice. Overall ANOVA *p* values and significant pairwise comparisons are indicated. **(D)** Corresponding gestational trajectories of the mean percentage of BrdU$^+$ cells within each lymphocyte population. Colored points and error bars represent group means ± SEM, and gray points represent individual mice. **(E)** BrdU$^+$ NK-cell event counts in the uterus/decidua, spleen, and PP before and across pregnancy. Overall Welch ANOVA *p* values and significant pairwise comparisons are shown. **(F)** Active caspase-3$^+$ NK-cell event counts in the decidua, spleen, and PP at GD7–8, GD12–14, and GD16–18. **(G, H)** Numbers of **(G)** BrdU$^+$ and **(H)** active caspase-3$^+$ cells within current T-bet-expressing TbetRFP$^+$NK1.1$^+$T-bet$^+$Eomes$^+$ cells, ex–T-bet TbetRFP$^+$NK1.1$^+$T-bet$^-$Eomes$^+$ cells, and non-fate-mapped TbetRFP$^-$NK1.1$^+$Eomes$^+$ cells across gestation. Bars represent means ± SEM, and each point represents one mouse. Brackets indicate comparisons with GD7–8. Significance is indicated in the figure. PP, Peyer's patches; FMO, fluorescence-minus-one.
